# Network variants are similar between task and rest states

**DOI:** 10.1101/2020.07.30.229492

**Authors:** Brian T. Kraus, Diana Perez, Zach Ladwig, Benjamin A. Seitzman, Ally Dworetsky, Steven E. Petersen, Caterina Gratton

**Author notes:** Correspondence: Caterina Gratton.

## Abstract

Recent work has demonstrated that individual-specific variations in functional networks (that we call “network variants”) can be identified in individuals using functional magnetic resonance imaging (fMRI). These network variants exhibit reliability over time with resting-state fMRI data. These properties have suggested that network variants may be trait-like markers of individual differences in brain organization. Another test of this conclusion would be to examine if network variants are stable between task and rest states. Here, we use precision data from the Midnight Scan Club (MSC) to demonstrate that (1) task data can be used to identify network variants reliably, (2) these network variants show substantial spatial overlap with those observed in rest, although state-specific effects are present, (3) network variants assign to similar canonical functional networks in different states, and (4) single tasks or a combination of multiple tasks produce similar network variants to rest. Together, these findings further reinforce the trait-like nature of network variants and demonstrate the utility of using task data to define network variants.

## 1 Introduction

An important issue in contemporary cognitive neuroscience is how to best identify individual differences in human brain organization that may be relevant to individual differences in behavior. It has long been established that measuring functional brain networks via fMRI can identify regions with correlated activity (i.e., “functional connectivity”) that support common functions such as motor processing and cognitive control (Biswal et al., 1995; Dosenbach et al., 2007). This organization of human brain networks has been mapped in multiple large groups (Power et al., 2011; Yeo et al., 2011) and more recently has been extended to individual-level brain networks (Braga & Buckner, 2017; Finn et al., 2015; Gordon, Laumann, Gilmore, et al., 2017; Gratton et al., 2019; Greene et al., 2019; Kong et al., 2019; Laumann et al., 2015, 2016; Marek et al., 2018; Miranda-Dominguez et al., 2014; Mueller et al., 2013; Poldrack et al., 2015; Seitzman et al., 2019; Sylvester et al., 2020). These networks can be measured during tasks or during a resting-state, where participants are asked to lie in the scanner without performing any particular task.

While there are many commonalities in network organization across individuals, some locations show large variations (Braga & Buckner, 2017; Gordon, Laumann, Adeyemo, Gilmore, et al., 2017; Gordon, Laumann, Adeyemo, & Petersen, 2017; Gordon, Laumann, Gilmore, et al., 2017; Gratton et al., 2018; Kong et al., 2019; Mueller et al., 2013; Seitzman et al., 2019). This variability across individuals is consistently larger than that found within individuals, suggesting that these individual variations represent meaningful “trait-like” differences in brain organization (Gratton et al., 2018; Seitzman et al., 2019). Individual differences in network organization are even sufficiently large to identify individuals based on their functional connectivity profile alone (Finn et al., 2015; Miranda-Dominguez et al., 2014).

A recent paper investigated the “trait-like” characteristics of individual differences in brain networks in additional detail (Seitzman et al., 2019). The authors found that particular punctate locations showed very large individual differences in functional network organization relative to the group (with correlations r < 0.3), that they termed “network variants”. All individuals showed evidence of network variants, although particular variants differed in location, size, number, and network association across individuals. Importantly, the authors demonstrated that not only were network variants common, but they were also quite reliable within individuals over resting-state scans, given sufficient data (achieving r > .8 with 40 min. of data).

However, one outstanding question concerning the “trait-like” nature of network variants is whether they are stable across different states, such as during task performance. In order to have utility as biomarkers, ideally network variants would not depend on ongoing cognition, such as what a person is thinking about during a scan session (Gratton et al., 2018). To date, network variants have only been identified using resting state fMRI data (Seitzman et al., 2019), and it is unknown whether similar variants can also be identified using task fMRI data. Recent evidence suggests that functional connectivity as a whole is stable between states with only small observable task-dependent changes (Cole et al., 2014; Gratton et al., 2018; Krienen et al., 2014), although individually-specific task-dependent changes were somewhat larger (Gratton et al., 2018). Thus, we sought to test whether these findings would also generalize to network variants.

A related practical consideration is that most existing datasets do not have sufficient resting-state data to achieve high reliability at the individual level (most have 5-10 min. of data, where reliability is poor; (Elliott et al., 2019; Gordon, Laumann, Gilmore, et al., 2017; Laumann et al., 2015; Noble et al., 2017)). However, many of these datasets have additional task data. Moreover, in populations where excessive motion is an issue, it can be difficult to collect resting state data whereas task fMRI may be more feasible (Greene et al., 2018; Vanderwal et al., 2019). One possibility is to combine rest and task data together to achieve higher reliability for network variants (as has been suggested by Elliot et al. (2019) for the connectome as a whole), which seems like a promising approach. However, for this approach to be effective, it is important to understand the degree to which the specific network properties under investigation are state dependent. While Gratton et al. (2018) demonstrated that task effects on functional connectivity as a whole were modest, they were present and significant. Thus, an important question for practical identification of network variants in other datasets, especially those of clinical interest, is to what degree their locations and network properties are state-dependent and can be computed from task-based measures instead. If task data can be used to identify similar network variants as observed in resting state data, this would make many more datasets which contain large amounts of task data suitable for this individual-specific analysis.

To address these theoretical and practical questions, we utilized data from the Midnight Scan Club (MSC). The MSC dataset is well-suited to examine these issues as it includes data from 10 individuals across 10 different sessions with approximately 10.5 total hours of fMRI data across 4 task states and rest. With such a large amount of data for task and rest states, we can compare (1) reliability of network variants identified using task data, (2) similarity in the *locations* of network variants between task and rest states, (3) similarity of network variant *connectivity profiles* between task and rest states, and (4) whether network variants identified in individual tasks differ from each other. Jointly, these analyses suggest that network variants are largely trait-like and that task data can be reasonably used for their identification and analysis.

## 2 Methods

### 2.1 Overview

The ability to use task data to measure network variants was investigated using several methods. (1) In the first analysis, we quantified the reliability (within-state) of network variants identified with task data (Figure 2). Next, we examined the stability (between-state) of network variant locations during task and rest (Figure 4). Third, we measured the stability (between-state) of network affiliations for variants (Figure 5). (4) To examine whether network variants show greater stability than expected relative to other areas of cortex, we compared the stability of network variant locations to the rest of the vertices on the cortex (Figure 6). (5) Finally, we examined whether network variants showed task-specific patterns of deviation from group functional connectivity (Figure 7). These analyses quantify both the degree to which network variants show trait-like stability and the feasibility of using task data to identify network variants.

**Figure 1.**
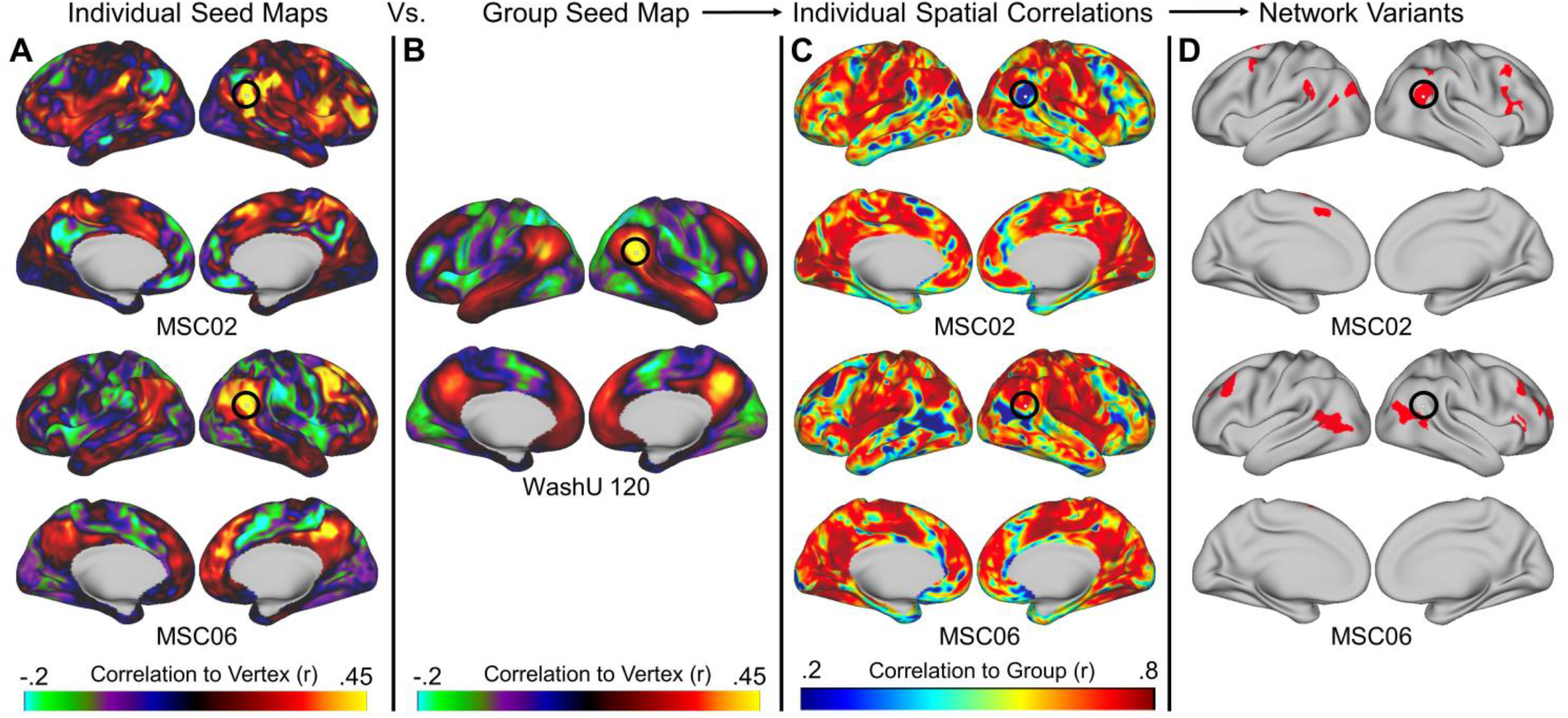
The process for generating network variants. First, a seed map is generated containing the correlations of every vertex to every other vertex within each individual during either task or rest (A) and the group average (B). Next, a spatial correlation between these maps is computed and the degree of similarity between each individual and the group is generated (C). Finally, regions with the lowest correlations (5%) with the group are selected as variants (D; variants are shown in red and small variants and locations in low signal regions are excluded). The seed maps shown here are for the vertex inside the black circles. This vertex corresponds to a network variant in MSC02 but not in MSC06.

**Figure 2.**
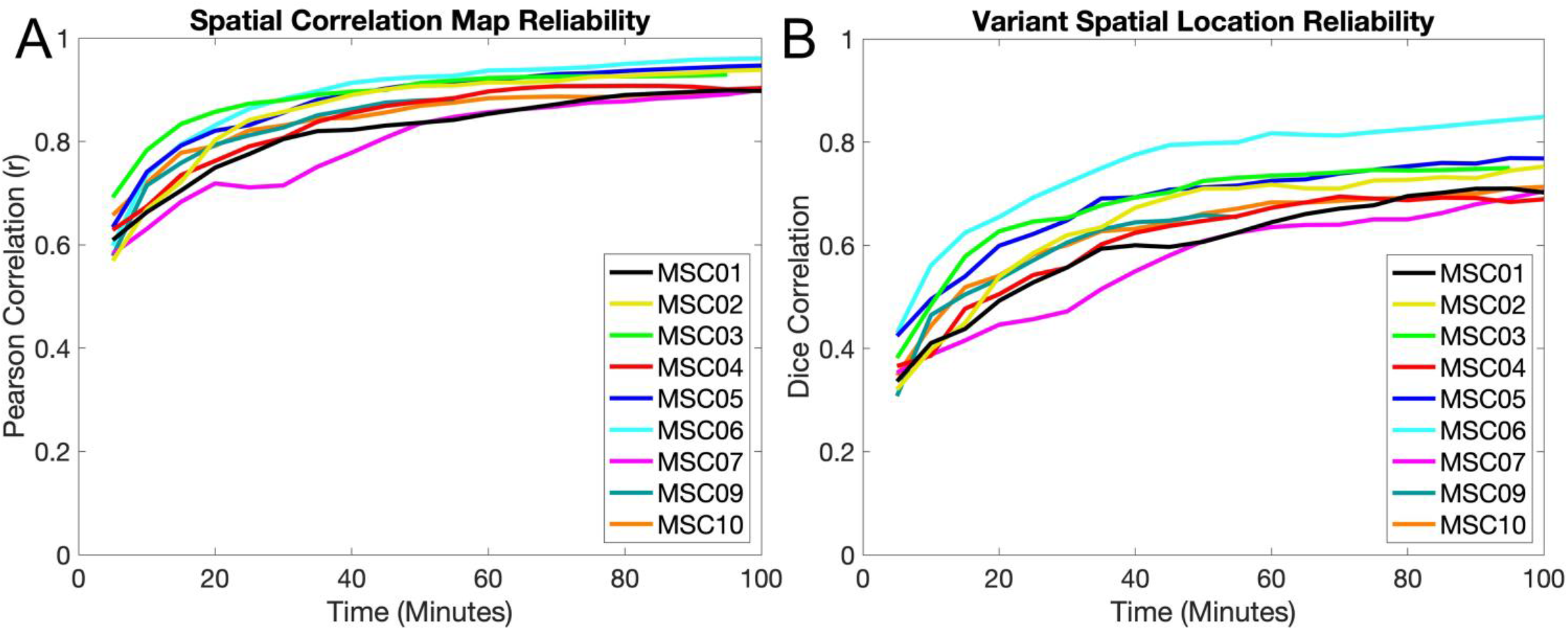
Reliability of network variants measured from task data. (A) Comparisons of individual-to-group spatial correlation maps from task data are shown for 5 minute increments for each participant. The similarity between maps is calculated via Pearson correlation. Spatial correlation maps begin to plateau around 30-40 minutes at r > 0.8. (B) Comparisons of network variant maps (after binarization of the spatial correlations to the bottom 5% of locations) from task data are shown in 5 minute increments for each participant. The similarity between binarized locations is calculated via Dice correlations. Binarized maps take longer (50-70 min.) and reach a slightly lower asymptote (Dice > 0.65), likely due to instability in the thresholding operation. Note that MSC09 (55 minutes) and MSC03 (95 minutes) did not have 100 minutes of data in their respective split-halves. A similar pattern was observed for the reliability of network variants in rest data (*Figure S1*). Similar analyses were also performed using the non-residualized task data (*Figure S2*) and data thresholded at r < .3 (*Figure S3*).

**Figure 3.**
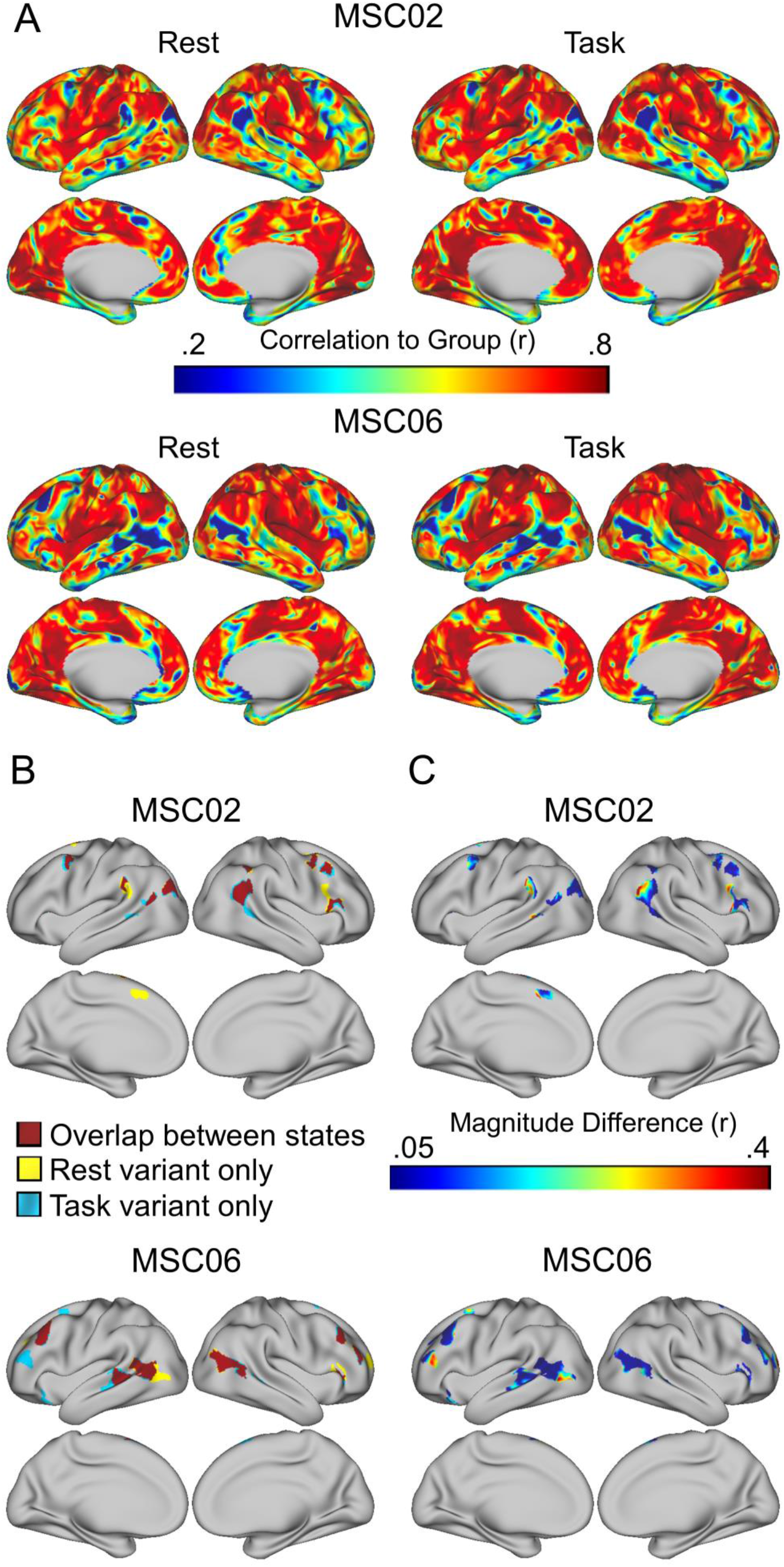
Comparison of network variant locations between task and rest states. (A) Individual-to-group spatial correlation maps are shown separately for task and rest states in MSC02 and MSC06; note the strong similarity in map profiles. (B) These maps are binarized (bottom 5%) to generate network variants for each state. The overlap between states is shown in red, and variant locations seen in only one state are shown in yellow (for rest) and blue (for task). (C) For each location labeled as a variant in either task or rest state, we have plotted the absolute difference in magnitude of the spatial correlations between states. Most locations show small magnitude differences, but a small number show more sizeable differences.

**Figure 4.**
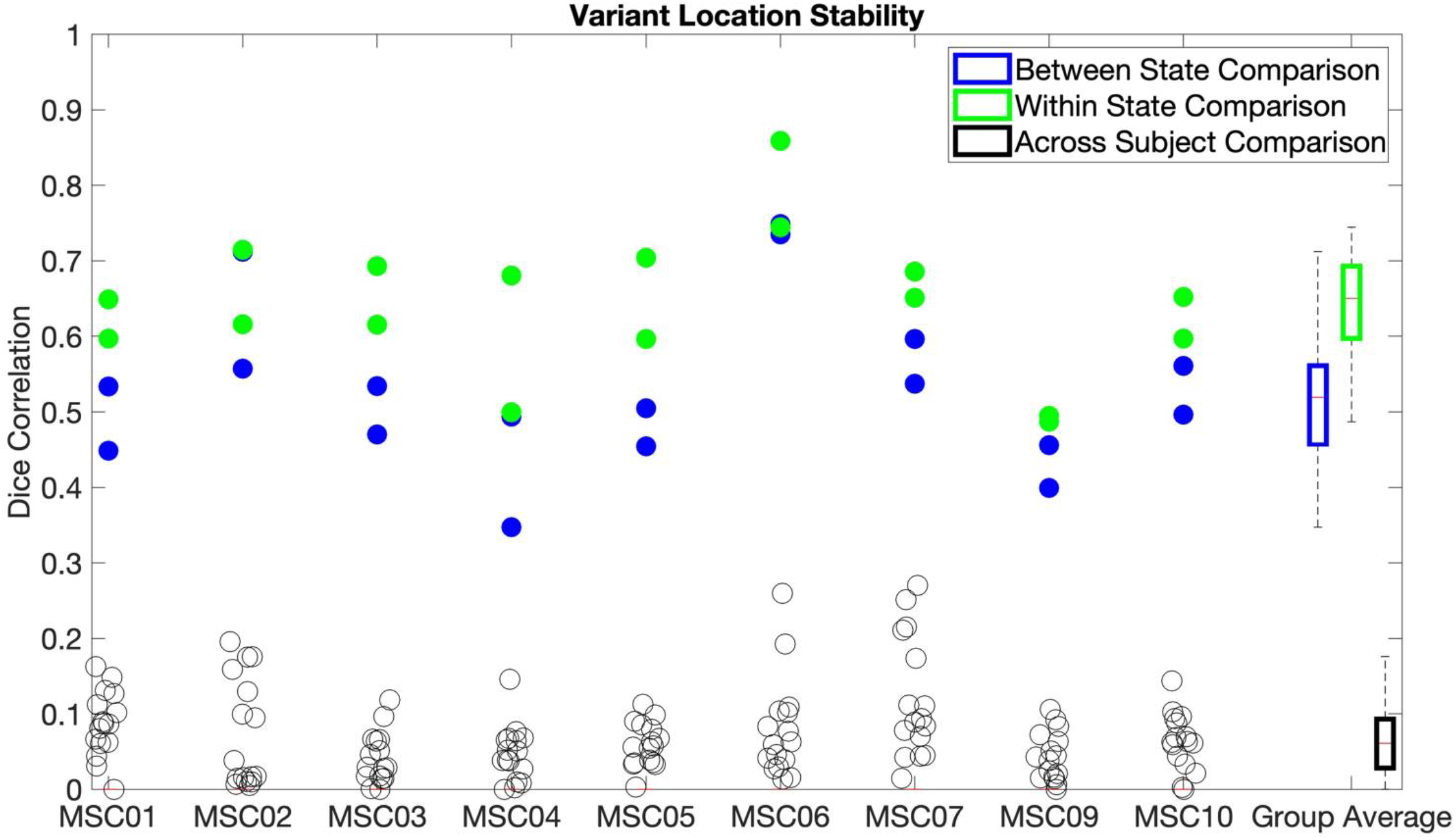
Network variant locations were compared using a Dice correlation, for network variants between states (blue), within states (green), and across subjects (black). Values are plotted for each subject (columns) with the summary boxplot at the end. Within and between state comparisons are shown as two dots, one for each pair of split-halves tested (see sections 2.9.2 and 2.9.3). The same results are also reported using a 2.5% threshold (*Figure S6*), for non-residualized data (*Figure S7*), and for data using an absolute threshold (r < .3; *Figure S8*).

**Figure 5.**
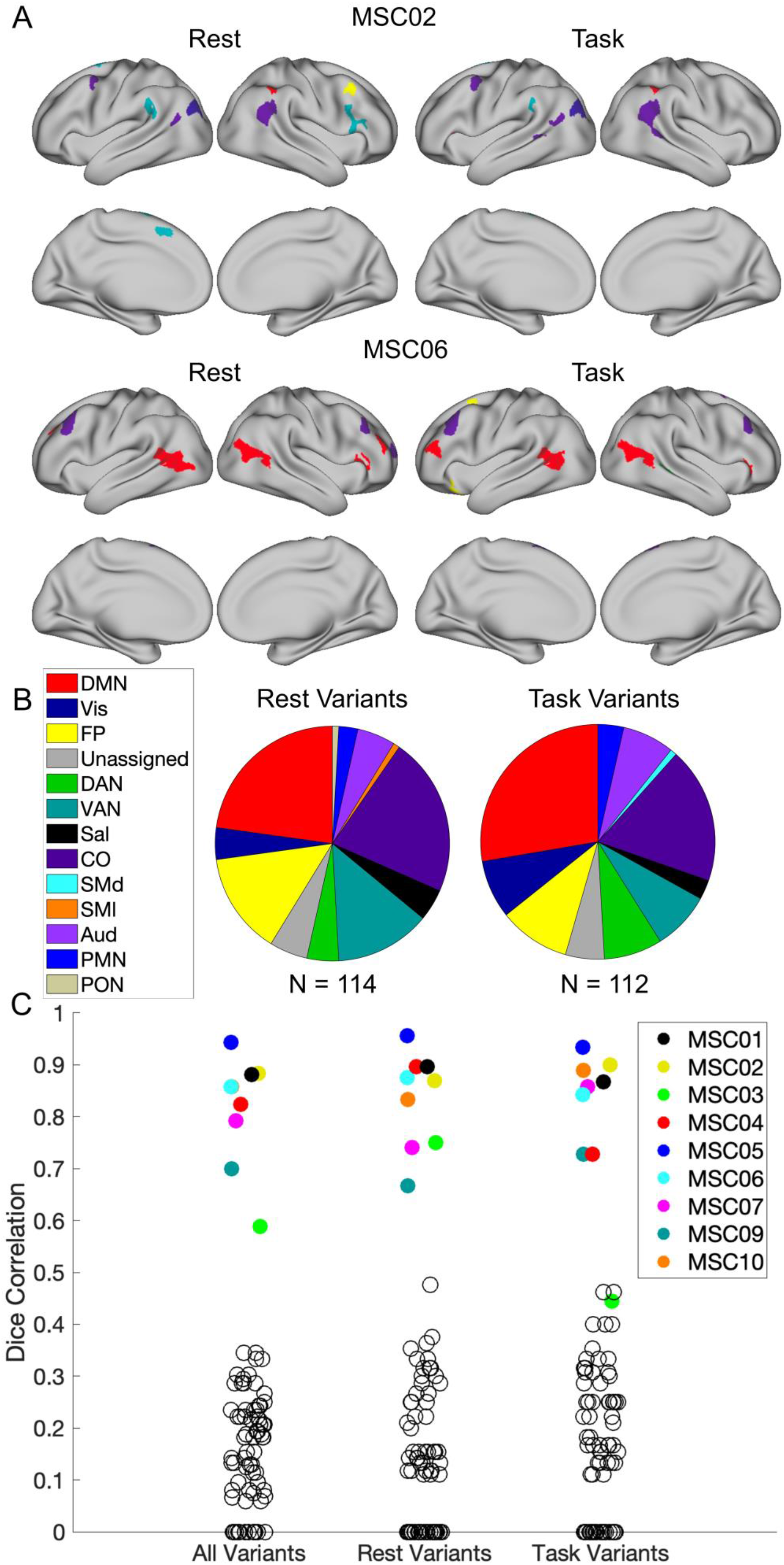
Comparison of network variant assignments between task and rest states. (A) The network assignments for variants are shown for task and rest states in MSC02 and MSC06. (B) The distributions of the networks that variants are assigned to for rest variants and task variants are also shown across all subjects. (C) The likelihood of a variant in one state being assigned to the same network in the opposite state is plotted for each participant. The empty black circles represent the likelihood of the variants in one subject being assigned to the same network in another subject. Comparisons were also made for whole-brain seedmaps (*Figure S10*) and for network assignments performed at the vertex-level rather than on contiguous variants (*Figure S11)*, where again strong consistency was seen across states relative to across people. *Note that MSC03 has far fewer variants than other participants in both states, and thus their data is much more susceptible to noise (see Figure S11 for evidence of high stability at the vertex level)*.

**Figure 6.**
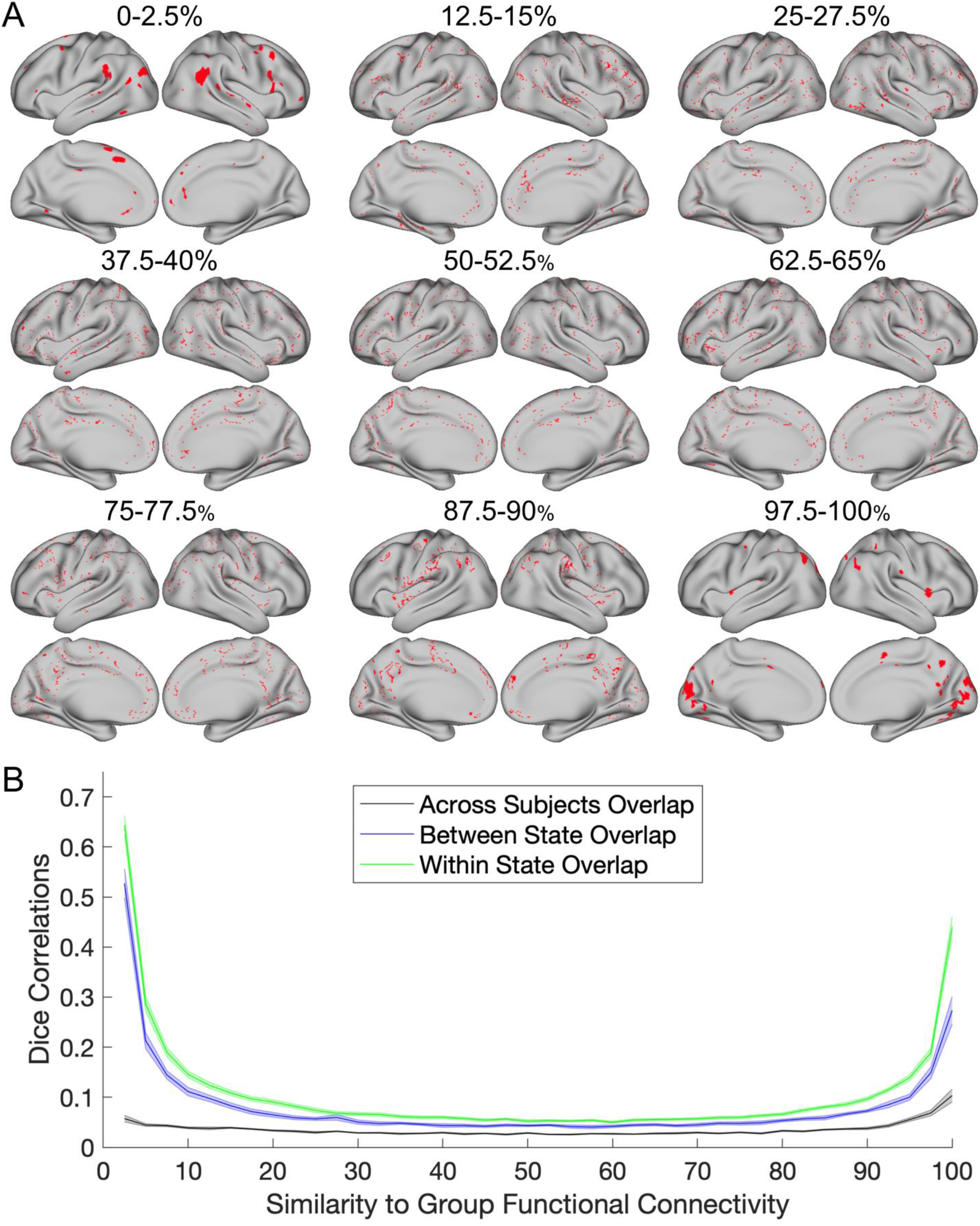
State/trait comparisons for locations binned by their similarity to the group (in 2.5% size bins). (A) The spatial distribution of the vertices in each bin are shown in red for MSC02. (B) At each bin, split-halves of the data were compared within state (green), between state (blue), and across subjects (black). Standard error bars are shown for each line. The areas that are the most similar and dissimilar to the group average functional connectivity show the most consistency and trait-like nature.

**Figure 7.**
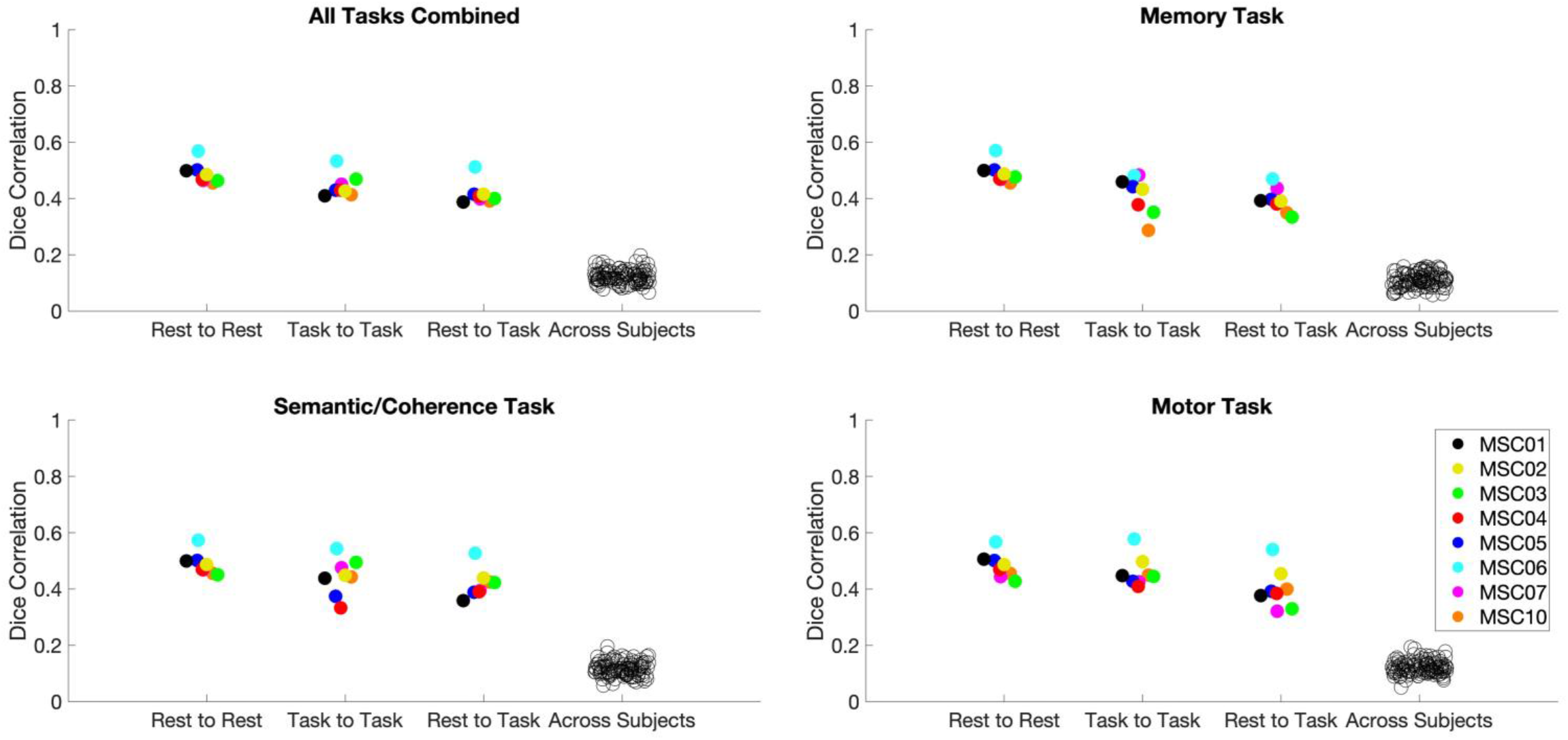
Comparison of network variant similarity in individual tasks relative to all tasks and rest. Dice correlations were used to compare network variants from data from all 3 tasks combined (upper left), the memory task (upper right), the semantic/coherence tasks (bottom left), and the motor task (bottom right). Each comparison shows the rest-to-rest, task-to-task (for the relevant task), rest-to-task (for the relevant task), and across subject comparisons of network variants. All comparisons were matched on amount of data within and across participants. Note that MSC09 was excluded from this analysis due to low amounts of high quality data in the motor task.

### 2.2 Datasets

The primary data used in this manuscript are from the Midnight Scan Club (MSC) dataset (Gordon, Laumann, Gilmore, et al., 2017). This dataset contains 5 hours of resting state data and approximately 5.5 hours of task data across 3 tasks (semantic/dot coherence, motor, and implicit memory) for 10 participants, collected across 10 separate fMRI sessions. One participant was excluded from analysis due to excessive head motion and drowsiness (Gordon, Laumann, Gilmore, et al., 2017; Laumann et al., 2016). A separate secondary dataset comprised of 120 neurotypical adults was used as the “group average” reference and is referred to as the WashU 120 (Power et al., 2013). These datasets are described in more detail elsewhere (Gordon, Laumann, Gilmore, et al., 2017; Power et al., 2013). The MSC and WashU MRI parameters are described briefly below. All data collection was approved by the Washington University Internal Review Board.

### 2.3 MRI Parameters

#### 2.3.1 MSC

The acquisition parameters for the MSC dataset have been fully described elsewhere (Gordon, Laumann, Gilmore, et al., 2017). In brief high-resolution T1-weighted (224 slices, isotropic 0.8 mm^3^ voxels, TE=3.74ms, TR=2400ms, TI=1000ms, flip angle=8 degrees), T2-weighted, and resting-state BOLD data were collected on a Siemens 3T Magnetom Tim Trio with a 12-channel head coil (gradient-echo EPI sequence, isotropic 4 mm^3^ voxels, TE=27ms, and TR=2200ms, whole brain acquisition).

#### 2.3.2 WashU 120

The acquisition parameters for this dataset have been fully described elsewhere. In brief, high-resolution T1-weighted (176 slices, isotropic 1 mm^3^ voxels, TE=3.06ms, TR=2400ms, TI=1000ms, flip angle=8 degrees), and resting-state BOLD data were collected on a Siemens 3T Magnetom Tim Trio with a 12-channel head coil (gradient-echo EPI sequence, isotropic 4 mm^3^ voxels, TE=27ms, and TR=2500ms). This dataset contains one 5 minute session of eyes-open resting state data per participant.

### 2.4 Task Designs and Analysis

The MSC dataset includes five different conditions. The WashU 120 contains only resting state data. The conditions in the MSC dataset are described in more detail elsewhere (Gordon, Laumann, Gilmore, et al., 2017; Gratton et al., 2018), but are briefly outlined below.

#### 2.4.1 Resting State

In the MSC, each session began with a 30-minute scan where participants were instructed to focus on a white fixation superimposed on a black background.

#### 2.4.2 Task Data

The MSC dataset contained data from four different tasks presented in counterbalanced order. Briefly, the motor task included two runs in each session (15.6 minutes total) of a blocked motor task adapted from the Human Connectome Project (Barch et al., 2013) where participants were cued to move either hands, feet, or tongue. The semantic and coherence tasks were presented as separate counter-balanced blocks in the same run two times in each session (14.2 minutes total). In the semantic task, participants responded to whether each word was a verb or a noun (both presented at 50% frequency). In the coherence task, Glass patterns were presented as white dots on a black screen (Glass, 1969). These dots varied how concentrically they were arranged, with either a 50% or 0% coherence to a concentric arrangement (both presented at 50% frequency). The semantic and coherence tasks were analyzed together as a mixed block-event related analysis (Gordon, Laumann, Gilmore, et al., 2017) and are referred to collectively as the “semantic/coherence” task in this manuscript. The memory task included three runs in each session (∼15 minutes total) with a different stimulus type (faces, scenes, words) presented in each run. In each run, participants were presented with 24 images, each of which was presented 3 times. Participants were asked to indicate whether the faces were male or female, the scenes were indoor or outdoor, and whether the words were abstract or concrete. Task activations were not directly analyzed in this manuscript, but task-related activity was calculated and regressed out from the timeseries via a general linear model as reported in Gratton et al. (2018).

### 2.5 Data and Code Availability

All of the data have been made publicly available (MSC and code: https://openneuro.org/datasets/ds000224/versions/00002; WashU 120: https://legacy.openfmri.org/dataset/ds000243/). Code for analysis related to network variants in MATLAB is available at: https://github.com/GrattonLab/SeitzmanGratton-2019-PNAS; other code related to MSC processing can be found at: https://github.com/MidnightScanClub. Code related specifically to the analyses in this paper will be located at this link upon publication: https://github.com/GrattonLab/.

### 2.6 MRI Processing

Data processing for the MSC dataset (Gordon, Laumann, Gilmore, et al., 2017) and WashU 120 (Power et al., 2013) are explained in detail elsewhere. The relevant details of the data processing are outlined below.

#### 2.6.1 Structural MRI Processing

For both datasets, the T1-weighted images for both datasets were processed via automatic segmentation of the white matter, grey matter, and ventricles in Freesurfer 5.3 (Fischl et al., 2002). The default recon-all command in Freesurfer was then applied to recreate the anatomical surface for each participant (Dale et al., 1999). In the MSC dataset, these surfaces were subsequently hand edited to improve the quality of registration. The surfaces were then registered to the fs_LR_32k surface space using the procedure described in Glasser et al. (2013). Using a separate calculation, a T1-to-Talairach transform was also performed (Talairach, 1988).

#### 2.6.2 Functional MRI Processing

The BOLD fMRI data from different runs in each session were concatenated within participants and pre-processed in the volume. First, a slice timing correction was applied, and the data were aligned to the first frame of the first run via rigid body transforms and then normalized to mode 1000 (Miezin et al., 2000). The functional data for the MSC dataset was then registered to the T2 image and subsequently to the T1 image which was previously registered to template space. In the WashU 120 dataset, the images were registered directly to the T1 image. Both datasets were then resampled to 3mm isotropic resolution and registered to the 711-2B atlas (Smith et al., 2004). For the MSC dataset, distortion correction was also applied in this step (Gordon, Laumann, Gilmore, et al., 2017). This correction was not applied to the WashU 120 dataset as no field maps were collected.

The task fMRI data were processed in the volume via a finite impulse response general linear model (GLM) approach as described elsewhere (Gratton et al., 2018). The residuals from the GLM were used to compute task functional connectivity using the “background connectivity” approach (Al-Aidroos et al., 2012; Fair et al., 2007). Subsequently, task residual processing was identical to the resting state data (i.e., completing all steps in the following section). As a supplemental analysis to assess whether regressing out the task activations affected state dependency, we also performed a few analyses without removing task activations from the data (referred to as non-residualized data here). Unless otherwise noted all references to task data in this paper were performed on the residuals from the task data.

#### 2.6.3 Functional Connectivity Processing

In the volume, additional preprocessing steps were applied to the data to remove artifacts from the data as outlined elsewhere (Gordon, Laumann, Gilmore, et al., 2017; Power et al., 2014). These steps included (1) the removal of nuisance signals via regression from the white matter, ventricles, global signal, motion parameters, as well as their derivatives, (2) removal of frames with high motion (FD > .2 mm, along with sequences containing less than 5 contiguous low motion frames, the first 30 seconds of each run, and runs with < 50 low motion frames (Power et al., 2014), and (3) bandpass filtering of the data (.009 Hz to .08 Hz). For MSC03 and MSC10, the motion parameters were low-pass filtered (below .1 Hz) before FD calculations to address respiratory activity in the motion traces (Fair et al., 2020; Gordon, Laumann, Gilmore, et al., 2017; Gratton et al., 2018, 2020; Laumann et al., 2016). After this preprocessing, the cortical functional data were registered to the surface. The cortical surfaces were transformed into the CIFTI data format in Connectome Workbench (Marcus et al., 2011). Lastly, a geodesic Gaussian smoothing kernel was applied (FWHM = 6mm, sigma = 2.55) using 2-D smoothing on the cortical surface.

### 2.7 Network Variants

#### 2.7.1 Functional Connectivity: Creation of Temporal Correlation Matrices

Correlation matrices were created for each participant for task and rest data separately based on timeseries correlations after censoring high motion frames. The amount of data (number of timepoints) used for correlations was matched in length across participants and states, approximately equally sampling across sessions (see *Table S1* for more details). The only exception to equally sampling across sessions was made for the case of the reliability analyses (section 3.1) where data was sampled consecutively from the first session until the total amount of required data (100 minutes) was reached rather than sampling an equal amount of data from each session. This sampling was selected for reliability estimates to provide a more ecologically valid assessment of reliability.

Based on these matched timeseries, a pairwise temporal correlation matrix was calculated for each participant between every pair of cortical surface vertices (59412 x 59412). The correlation matrix was then Fisher transformed. Thus, each row of this matrix represents a seed map for a given seed (vertex) to all other vertices on the surface (Figure 1A; (Seitzman et al., 2019). We also created a similar correlation matrix for the WashU 120, but in this case, the individual subject correlation matrices were averaged together to create a single group average matrix (Figure 1B).

#### 2.7.2 Spatial Correlations to the Group Average

In order to identify regions with strong dissimilarity from the typical functional connectivity profile (i.e., network variants), each MSC participant’s correlation matrix was contrasted with the group average correlation matrix from the WashU 120 dataset using a spatial correlation. To compute the correlation, the seed map for each vertex (i.e., a row of the correlation matrix) in a given individual was correlated with the corresponding vertex’s seed map in the WashU 120 group data, yielding one spatial correlation value per vertex (Figure 1C). This was repeated for each individual in the MSC (Seitzman et al., 2019). Vertices with low signal (mean BOLD signal < 750 after mode 1000 normalization) were masked (Gordon et al., 2016; Ojemann et al., 1997).

#### 2.7.3 Calculation of Network Variants

Next, in order to select regions that were most different from the group (“network variants”), the spatial correlation maps were binarized to keep only the lowest 5% of correlations to the group for each participant (values in the lowest 5% were set to 1, all other vertices were set to 0; Figure 1D; note this threshold differs slightly from Seitzman et al. (2019) due to small changes in the processing stream of the MSC data). Variants composed of less than 50 contiguous vertices and vertices with an SNR < 750 were excluded from analysis (Seitzman et al., 2019). The remaining vertices in the lowest 5% of correlations were then considered for further analysis. This procedure was used to create network variants in both task and rest data separately. To confirm the findings were not dependent on threshold, the analyses reported here are also shown at a 2.5% threshold in the *Supplementary Results*. For the 2.5% threshold, variants were required to be larger than 25 contiguous vertices. In addition, we also performed several analyses using an absolute threshold (r < .3) rather than a relative threshold. For these analyses, variants of less than 50 contiguous vertices were excluded from analysis.

### 2.8 Within-state Reliability of Network Variants in Task Data

If task data is to be considered a suitable substitute for rest in defining network variants, it must show similar within-state reliability as rest (as well as similar topography and network properties – tackled in the next sub-sections). To test how much task data is necessary to achieve reliable estimates of functional connectivity within an individual, each participant’s data was divided into split-halves composed of either odd or even numbered sessions. As above, data was equally sampled across all sessions for task and rest. Data from all 3 tasks was used in the task analyses unless otherwise noted.

Following Laumann (2015) and Gordon (2017), functional connectivity for each even numbered session was estimated using all of the available task data and this split-half was treated as the best estimate of “true” functional connectivity. In each odd numbered session, data was consecutively sampled starting with the first session in 5 minute increments (Seitzman et al., 2019). When all of the task data in a session was exhausted, data was then sampled from the next session until either 100 minutes of task data was sampled, or no more task data was available (see *Table S1* for details on the data sampling of each analysis). The data from these sessions was used as a test split-half.

For each functional connectivity matrix in both segments of data, a spatial correlation map versus a group average was computed using the steps outlined above. The similarity of the individual-to-group spatial correlation maps was then compared between the “true” half and progressive amounts of data from the test split-half via a Pearson correlation. Network variants were created using the procedure outlined in section 2.7.3, only no exclusion for variant size was applied for any of these analyses. These binarized maps were compared between the “true” and test halves via a Dice-Sorenson correlation (Dice, 1945; Sorensen, 1948).

### 2.9 Comparing Variant Spatial Locations

#### 2.9.1 Comparison of Variant Locations

To inspect the similarity of network variant locations across task and rest states, spatial overlap maps were created for each participant. These maps were created by determining which vertices on the surface were considered variants in both task and rest states within each participant. To quantify the amount of spatial overlap between states, variants were separately defined for rest and task states in each split half resulting in 4 variant maps (35.2 minutes total for each split-half matrix per participant; see *Table S1 -* data was approximately equally sampled across sessions in task and rest). Then, a Dice correlation was calculated to determine the similarity in variant locations within and between states for each participant.

#### 2.9.2 Between State Comparison

To compare the spatial overlap of variants between task and rest, each split-half of rest data was separately compared to each split-half of task data. This process was repeated for both pairs of split-halves (odd rest split-half to even task split-half and even rest split-half to odd task split-half) to produce two comparisons per participant. The overlap of the split-halves between states was computed via a Dice correlation. Paired t-tests were performed using the averaged value of the between-state comparisons within each participant.

#### 2.9.3 Within State Comparison

To compare the amount of spatial overlap of variants within task and rest, the split-halves of rest and task were both separately compared (odd rest split-half to even rest split-half and odd task split-half to even task split-half) to produce two comparisons per participant. The overlap of the split-halves within states was computed via a Dice correlation. Paired t-tests were reported using the averaged values from the within each state comparisons within each participant.

#### 2.9.4 Across Subject Comparison

To better evaluate the relative similarity of variants within participants (within and between states), we created a comparison benchmark of the similarity of variant locations across subjects. A given participant’s variants (computed from a single split-half) were compared to the variants from other participants (also computed from a single split half, from the opposite state). For instance, one split-half of the task data for MSC01 was compared with the corresponding split-half of rest data for MSC02. The similarity in variant maps was computed with a Dice correlation. This was repeated for all combinations of subjects and split-halves, yielding 128 comparisons. This benchmark described the likelihood of observing variants in the same location across subjects which could be used as a comparison for the likelihood of observing variants in the same location within subjects (either within or between states). To compute t-statistics for this comparison, all of the comparisons for each participant’s data were averaged into one value for that subject.

#### 2.9.5 Comparison of Variant Magnitude Differences Across States

To compare the magnitude of individual differences between states, variants were defined using the procedure outlined in section 2.7.3. Then, the spatial correlation magnitude at each of these vertices was calculated in both states. Finally, the mean absolute difference between rest and task magnitudes was calculated.

To test whether these values were bigger than what would be expected in other areas of the cortex, the variants in each participant were randomly rotated, thereby matching for variant size and shape in the null distribution (Gordon, Laumann, Adeyemo, Gilmore, et al., 2017). Within each participant, 1000 rotations were randomly generated and performed within each hemisphere. In the case that a rotation resulted in parcels that overlapped with the medial wall, this rotation was recalculated until none of the rotated parcels overlapped with each other or the medial wall. For each of the 1000 rotations, the mean absolute difference between task and rest magnitudes was calculated. These values were then compared to the mean absolute difference of variant locations in each participant.

### 2.10 Network Variants Versus Other Vertices

An outstanding issue in the measurement of network variants is whether they are more similar between states than would be expected from regions with more similarity to the group. Such a finding would provide evidence for their trait-like stability, as posited elsewhere (Seitzman et al., 2019). In order to examine this issue, we thresholded maps at 2.5% increments starting with the lowest 2.5% of spatial correlation values versus the group average for each split-half within each participant (e.g., bin 1: 0-2.5% similarity to the group, bin 2: 2.5-5%, bin 3: 5-7.5%, …, etc.). For each 2.5% increment, we computed the Dice correlation for the comparisons within-state, between-state, and across-subjects. The average value for each of these comparisons was then plotted with their associated standard error (across subjects). No size exclusion based on a minimum number of contiguous vertices was applied to this analysis.

### 2.11 Similarity of Individual Tasks Versus Multiple Tasks

We also evaluated the degree to which variants at rest are similar to variants from a single task versus variants from multiple tasks averaged together. As above, the amount of data was matched across individual tasks and participants (see *Table S1*). Data were sampled into split-halves using the same sampling procedure as outlined above. This same procedure was also applied to sample an equal amount of data from all three tasks combined. The amount of data for each individual task and for all three tasks combined was 11.3 minutes total per split-half for each task per participant. MSC09 was not used in this analysis due to the small amount available for this participant (1.25 minutes in the smaller split-half). Dice correlations were used to quantify the overlap within-state, between-state, and across-subjects. Within subject ANOVAs were used to test for differences between states. Due to the small amount of data available for this analysis, the results should be interpreted with some caution.

### 2.12 Similarity of network variant assignments between states

We next asked whether variants also showed similar network assignments between states. To test this, each contiguous variant unit was assigned to a network, and the similarity of these networks assignments between states was evaluated.

#### 2.12.1 Assigning Network Variants to Functional Networks

Each variant was assigned to functional networks via a winner-take-all algorithm. This algorithm assigned variants to one of 14 group average template networks (Gordon, Laumann, Adeyemo, Gilmore, et al., 2017). The templates for matching the network variants were the spatial maps of the group average canonical networks from the group average of the WashU 120 dataset. First, a seed map was generated for every vertex composing a given variant. These seed maps were then averaged together to form one seed map for each network variant (Seitzman et al., 2019). For each map and the template, the highest 5% of correlations with the rest of the surface were binarized. Next, we computed the similarity (Dice correlation) between the binarized network variant map and the templates. Vertices within 30mm geodesic distance from any vertex in each variant were excluded from the variant seed map and template seed map (Gordon, Laumann, Adeyemo, Gilmore, et al., 2017). Each network variant was assigned to the winning template with the largest Dice correlation. If no good “winning” template was found based on the lowest 5% of Dice correlations with the winning network (Dice < 0.1428 at 5% threshold) or multiple networks tied for the highest value, then the variant was assigned to an “unknown” network. In addition, network variants were also removed if their assigned network spatially overlapped at least 50% with the group average network (9/123 = 7% in rest, 11/123 = 9% in task; (Seitzman et al., 2019).

#### 2.12.2 Evaluating Variant Network Assignment Consistency Between States

We next asked whether a particular variant showed the same network assignments between states. To do this, first variants in task and rest were assigned to networks according to the procedure above. Next, the same vertices were selected in the other state (e.g., the variant locations from a task were applied to the resting state data), and the network assignment procedure was repeated for these vertices in the other state. A Dice correlation was then used to quantify the likelihood of the same vertices between states being assigned to the same network (with matching variants set to 1 and non-matches set to 0). To test whether variant assignment was more stable between states than across subjects, we also used this method to quantify how often the same vertices assigned to the same network in all other subjects.

Additionally, we also examined whether assigning contiguous variant units to networks produced different results versus assigning each individual vertex composing a variant to a network (given that contiguous variant units may not be homogenous). The network assignment procedure was identical, but instead of using the average seedmap for each variant, the seedmap for each vertex composing each variant was assigned to a network. We then compared the distributions vertices assigned to each network and the similarity of vertex-level assignments across states.

We also examined the consistency of network variants’ network patterns between states by comparing the correlation of the full seedmap across states. For each variant, a seedmap was calculated for each vertex and then all of the seedmaps contained in a variant were averaged together. This same process was then performed for that variant in the opposite state. As a comparison, this was also repeated for the same location in the opposite state across all the other subjects. The similarity of these maps was computed via a spatial correlation.

### 2.13 Relationship Between Task Activations and Functional Connectivity

In an additional analysis, we explored whether the amount of task activation relative to baseline was associated with the likelihood of a variant showing inconsistent patterns across task states. If task activations systematically alter functional connectivity, then it could be that areas which show more task activations may be identified as variants in task states but not during rest. To test this question, the average activation was calculated separately for vertices which were identified as variants in both task and rest states (overlapping vertices), vertices which were identified as variants only in rest states, and variants which were identified as variants only in task states (see Figure 3B). The average task activation for each subject was then calculated for variants identified in both states (overlapping), task states, and rest states. Each parcel of contiguous vertices in each state had to be composed of at least 30 vertices to be included in this analysis. These values were compared via a t-test.

## 3 Results

### 3.1 High Amounts of Task Data Can Produce Reliable Estimates of Network Variants

The overarching question of this manuscript is whether network variants are similar between task and rest states, indicating that they are trait-like and that task data may be used to identify variants. As a first step, we asked if task data can produce reliable (i.e., high within-state similarity) estimates of network variants, as has been shown for rest data. Following Gordon et al. (2017) and Laumann et al. (2015), reliability was estimated using a split-half procedure. Specifically, task data from each participant was divided into two halves. Variants were created from one half of the data (mean = 109 minutes, range = 75.9-123.9 minutes); these variants were treated as our best estimate of “true” variants. The other half was incrementally sampled in 5 minute increments of data; each incremental sample was used to generate a new estimate of network variants. This incremental sample estimate was then compared to the other full half (the “true” estimate). The same procedure was conducted both for the comparisons of the continuous spatial correlation maps (Figure 2A, compared using Pearson correlations) and for the binarized network variants (Figure 2B, compared using Dice correlations).

The general trend is that as the data quantity increases (x-axis), the similarity between the two halves of data also increases (y-axis) before reaching an asymptote, typically around 40 min. of data. In particular, with > 40 minutes the continuous spatial correlation maps reach reliabilities with r > .8 (Figure 2A). In addition, the binary overlap between the lowest 5% of correlations between the group average also becomes stable with high amounts of data (Figure 2B; note that however these values typically asymptote at a lower level, around Dice correlation = 0.7, likely due to instability caused by the thresholding operation). However, reaching stability for the binarized data requires additional time beyond what is necessary to achieve reliable spatial correlation maps (> 70 minutes). This task reliability profile is similar to what is seen with rest (*Figure S1*). Similar results are also obtained using non-residualized task data (i.e., data without task activation removal, treated the same as rest; *Figure S2*) and when using an absolute (r < .3) rather than percentile threshold (*Figure S3*; though the results are more variable across subjects than when a percent threshold is used). Thus, high amounts of task data can produce reliable measures of network deviation in individuals with respect to the group average, similar to rest.

### 3.2 Network Variants Occur in Relatively Similar Locations Across States

Next, we examined the similarity of network variants across states. The number of variants (*Table S2*) and vertices (when thresholded at r<0.3; *Table S3*) in both task and rest states are summarized in the supplemental tables. We did not see significant differences in variant quantity across states (all *p* > 0.05).

We next focused on whether network variants occur in the same locations between task and rest. If network variants are trait-like, then their vertices would be expected to overlap at a significantly greater rate between states (within a person) than across people. Consistent with this pattern, Figure 3A,B shows that network variants appear in similar locations between task and rest states in single individuals (see *Figure S4* for variant overlap for the other participants).

To quantify this similarity, we calculated the Dice coefficient between task and rest state network variants. We benchmarked this between-state overlap against two comparisons: (a) within-state overlap (how similar network variants are across two different samples from the same state, e.g., between two split-halves of resting state data from MSC01) and (b) across-subject overlap (how similar network variants are between subjects, e.g., between a task split-half of MSC01 and a rest split-half from MSC02). These comparisons allowed us to determine if network variants were more trait-like (showing relatively similar variants between states, but distinct variants across people) or state-like (showing relatively distinct variants between states, closer to the level seen across people). Importantly, data quantities were matched for all comparison samples within and across individuals.

The results showed that variants were more likely to overlap between states (*M* = .533, *SD* = .103) than across subjects (*M* = .068, *SD* = .025, *t*(8) = 14.96, *p* < .0001, *d* = 4.99), suggesting a large trait-like component to network variants. Variants were also significantly more likely to overlap within states (*M* = .641, *SD* = .082) than between states (*M* = .533, *SD* = .103, *t*(8) = 6.38, *p* = .0002, *d* = 2.13; see Figure 4), suggesting some state-dependence. However, the magnitude of the difference between states was substantially smaller than the magnitude of the difference across subjects. Similar results are also observed at a 2.5% threshold (see *Figure S5*) as well as when using an absolute (r < .3) rather than percentile threshold (see *Figure S6*). Non-residualized task data (where task activations were not removed) showed a similar overall pattern, but significantly lower between-state stability (*t*(8) = 8.09, *p* < .0001, *d* = 2.7; *Figure S7*), suggesting that it may be important to address task activations to improve network variant correspondence between task and rest. Jointly, these findings indicate that variant locations are affected by state, but are relatively more trait-like than state-like. We also examined whether the similarity to the group map differed at network variant locations more than for other areas of the cortex (Figure 3C). On average, the differences between states are relatively small for network variants (*M* = .102, *SD* = .016, *Skew* = .824). However, the tail of the distribution for magnitude differences is somewhat larger for network variants than would be expected by chance in other areas of the cortex (*M* = .064, *SD* = .012, *Skew* = .522), indicating that a subset of locations shows larger magnitude differences than might be expected for other areas of cortex (see *Figure S8*). This is consistent with some locations showing an interaction between individual differences in functional connectivity and task state.

Finally, we asked whether task activations were related to inconsistencies in network variant identification between states. To examine this, each variant was divided up into sections depending on whether its vertices were identified as variants in both task and rest states (overlapping), in rest only, or in task only. The average task activation was then calculated separately for each variant section (*Figure S9*). The mean task activations did not differ by variant sub-units for any of the tasks (*p* > .05). This suggests that state-dependent differences in network variants are not simply driven by average task activations. Future studies will be needed to examine if more nuanced relationships link specific activations to specific network variants.

### 3.3 Network Variants Associate with Similar Networks in Task and Rest

While the locations of network variants are relatively similar across states, it is possible that the variants may show different functional correlation maps, matching to different networks across states. To test this possibility, each variant was assigned to a template network using a winner-take-all procedure in each state (see section 2.12.1). A Dice correlation was then used to quantify how often variants were assigned to the same network between states. As shown in Figure 5B, the distribution of variant network assignments between states was similar. In addition, the likelihood of obtaining a matching network assignment for a variant using the same vertices between states was high (*M* = .814, *SD* = .109; All Variants in Figure 5C). This was significantly greater than the likelihood of variants matching across subjects (*M* = .159, *SD* = .056), *t*(8) = 20.03, *p* <. 0001, *d* = 6.68. This shows that network variants tend to be assigned to the same network between states, providing evidence for their trait-like stability.

We also looked at the similarity of seedmaps for each variant (rather than using a winner-take-all procedure to assign to one of 14 networks). The purpose of this analysis was to examine the similarity of seedmaps between states and across subjects while avoiding the non-linearities inherent in the network assignment procedure. Variant seedmaps showed a very high similarity between states (*M* = .824, *SD* = .033), much higher than across subjects (*M* = .034, *SD* = .041), *t*(8) = 33.07, *p* <. 0001, *d* = 11.02 (see *Figure S10*), suggesting that the network profiles of variant regions are similar between states.

The analyses above assume that contiguous variants represent single homogenous regions, with the same network assignment, which may not always be the case distorting differences in network assignment across states. To test if treating contiguous variants as a single unit contributes to state-level differences in variants’ network assignments, we assigned single variant vertices to a network and repeated the procedure (*Figure S11A*). The distribution of the network assignments was extremely similar across states and to that seen with contiguous unit assignment. Additionally, variant vertices were much more likely to assign to the same network between states (*M* = .819, *SD* = .031) than across subjects (*M* = .193, *SD* = .028), *t*(8) = 41.69, *p* <. 0001, *d* = 13.9 (*Figure S11B*). These results confirm that variants show similar network profiles between task and rest states, whether they are considered homogenous units or as individual vertices.

### 3.4 Network Variants are Highly Stable Compared to Vertices More Similar to the Group

Next, we looked at whether network variants are unique trait-like clusters relative to other subsets of vertices selected based on their similarity to the group. To test this, we binned vertices into sub-units based on their similarity to a group average map, e.g., 2.5-5%, 5-7.5%, 7.5-10%, etc. most similar to the group.

The results of this analysis are shown in Figure 6. Interestingly, only bins on the extreme low and high ends of the curve contained vertices which formed contiguous clusters (Figure 6A). Thus, regions of functional connectivity that are most similar and most dissimilar to the group average tend to represent focal regions. Not surprisingly given this pattern, network variant locations (the left side of the curve; Figure 6B) show more stability between states, with higher within- and between-state similarity than almost any other percentile bin. The only other bins that showed as strong of stability were at the highest end of the spectrum: those with the most consistency to the group. This is likely driven by the fact that network variants (as well as the most group consistent regions) agglomerate into contiguous parcels, whereas other bins show a more scattered representation across cortex. This provides evidence for the distinctive nature of network variants relative to other locations selected for their group similarity.

### 3.5 Network Variant Stability was Similar When Examined for Single Tasks

The last comparison tested was whether variant stability differed between any of the 3 tasks collected in the MSC. A significant effect would indicate that there were task-specific effects of how stable variants were across states. To test this effect, similar overlap analyses as those presented in Figure 4 were carried out, but with data separated by task rather than concatenating across all tasks. Note that, given the separation, this analysis was conducted on relatively less data (11.3 minutes per split-half) and thus the results should be interpreted with caution due to lower reliability.

Overall, the results suggest that variant similarity between task and rest does not significantly differ substantially between tasks (Figure 7). To test whether the stability of network variants differed between individual task states, a one-way within subjects ANOVA was run on the values from the rest-to-task comparisons for each task. Greenhouse-Geisser corrections were applied where appropriate. The effect of task was not significant, (*F*(1.631,11.418) = 1.565, *p* = .248, *d* = .95), indicating that there were no significant differences in stability for any individual task versus rest. This suggests that there is no difference in the stability of network variants when using a single task versus multiple tasks compared to rest. Similar results are also seen when using a 2.5% threshold to identify network variants (see *Figure S13*).

## 4 Discussion

This study examined whether network variants (locations where an individual’s network organization differs markedly from the group) were susceptible to state-level changes induced by task demands. This question has bearing on both the trait-like aspects of network variants and on practical questions of whether task data can reasonably be used to identify network variants. To examine this question, we used resting state and task data from the Midnight Scan Club to define network variants. These network variants were then compared to each other in various ways to evaluate whether network variants showed trait-like stability between states. We found that (1) network variants defined from task data show similar reliability to resting state data, (2) network variants appear in similar locations between task and rest data, although with some small state-dependent effects, (3) the network assignments of variants are similar between task and rest, and (4) that there is similar stability for network variants identified from multiple or single tasks. Thus, our analyses suggest that while network variants show some differences that are due to state-level factors, they are predominantly stable between states, consistent with their trait-like properties. Importantly, these findings suggest that a reasonable approximation of network variants can be identified from task as well as rest.

### 4.1 Network Variants Are Largely Trait-like

The current findings add an additional piece of evidence to suggest that network variants are relatively trait-like in nature by demonstrating consistency across task states. We found that individuals’ network variants tend to occur in similar locations across different states as well as have similar network labels. These findings add to previous evidence from Seitzman et al. (2019) showing (a) that network variants are consistent across sessions within a state, even over a year, (b) that variants defined at rest can be used to predict default mode deactivations during a task and (c) that variants can be used to separate individuals into sub-groups that relate to behavioral measures taken outside of the scanner. This consistency across states, high reliability, and relationship to brain activation and behavioral measures taken at other moments in time jointly suggest that network variants may be trait-like properties of brain networks, rather than functions of ongoing processing.

Taken together, these results suggest that the neurobiological underpinnings of network variants may stem from a systematic change in the function of underlying brain tissue from that which is typically observed in the population, which then alters the Hebbian associations between brain regions and their functional connectivity properties (Gratton et al., 2019; Seitzman et al., 2019). Given their stability, these individual-level differences in brain network organization may be due to differences in cortical organization driven by longer-term factors such as genetics, prolonged life experiences, or other environmental factors. Under this framework, network variants may be viewed as prime candidates for neural correlates of trait-like individual differences in behavior, such as cognitive ability and risk for psychopathology.

Although network variants primarily exhibit trait-like properties, it is also clear that they show some minor but significant state dependence. This is consistent with past work showing that the functional organization of the brain is largely stable, but can show subtle modifications in adjusting to different task states (Cole et al., 2014; Gratton et al., 2016; Krienen et al., 2014). In the context of individual differences, recent work has emphasized that stable group and individual factors dominate brain networks, but more modest task-based effects are also present in functional organization between task and rest states across the full connectome (Gratton et al., 2018). These state-dependent changes in functional connectivity likely also contribute to differences in connectivity of network variants between different task states.

One possible explanation is that, if network variants are areas which have shifted in their functions from their ‘typical’ role, this may also compound differences during particular task states, dependent on how the task affects this atypical neuronal population and its coherence with other regions. While there is some evidence that functional connectivity changes more in task active regions (Gratton et al., 2016), we did not find that variant-dependent shifts were related to overall task activations (see *Fig. S9*). It is possible that variant state changes are related to more nuanced relationships between specific variant types and particular task processes. A full examination of this question will benefit from future studies adopting specialized tasks with contrasts targeting discrete networks (e.g.,(Braga et al., 2019; DiNicola et al., 2020)). Future studies with a larger sample of tasks and participants will also elucidate whether these state-dependent changes in variants are systematically related to unique strategies or task performance (performance on the current tasks, which were all relatively simple, was at ceiling and all MSC individuals are high IQ).

### 4.2 Practical Considerations for Identifying Network Variants With Task Data

Due to the similarities of network variants between task and rest states, it seems reasonable that network variants can be estimated using task data. Network variants show substantial stability in location and network characteristics between rest and task states (*M* = .631; see *Figure S13*). In practice, there is approximately a 17% reduction in the stability of network variant location when examined between states (*M* = .533) versus within states (*M* = .641). Correlation patterns and network assignments show an even higher stability between states (*M* = .814). While identifying variants between states is not as consistent as identifying them from within the same state, the results suggest that it is reasonable to conduct network variant analyses across tasks if needed for data quantity reasons. Notably, spatial differences in network variants between states are much smaller than the differences observed in reliability from using small amounts of data (approximately 56% lower between 5 minutes and 70 minutes of data).

Critically, the finding that variants can be identified during task states with good correspondence to rest opens up new datasets for variant analysis that would not be available otherwise. While most currently available datasets only have small amounts of rest data (5-10 mins) which are insufficient to achieve high reliability (Fig. 2B, S1B), many have substantial additional task data. Moreover, while datasets from clinical populations are of great interest, it may be particularly difficult to obtain large amounts of high quality resting state data in many of these populations (due to exacerbated head motion, compliance issues, etc. (Greene et al., 2018; Hodgson et al., 2017; Vanderwal et al., 2019)). Combining data across task and rest may be an effective strategy to increase functional connectivity reliability sufficient for network variant analyses without sacrificing substantial variability due to state differences. An important caveat to this is that accounting for task activations may be critical to ensuring higher similarity between rest and task states. Because of this, recent work suggesting movie watching (where it is harder to remove evoked signals) as an alternative to rest may not be ideal for combining data across states.

Elliot and colleagues (2019) have shown that combining resting state data with background connectivity during task performance increases the reliability of functional connectivity data as a whole. Critically, this increase in reliability was also accompanied by an increase in the predictive utility of functional connectivity for cognitive measures and the heritability of functional connectivity. Additional work has also incorporated this strategy of analyzing functional connectivity data in developmental samples (Cui et al., 2020), demonstrating its utility in a dataset with a limited amount of resting-state fMRI data. Our results suggest that this method can also be used to identify network variants, which may yield new insights into neurologic and psychiatric disease.

### 4.3 Important Methodological Considerations for Identifying Network Variants

Despite this promise for using task data to identify network variants, some caveats for this approach are discussed below. First, while the reliability of cortical functional connectivity overall becomes stable with >30 minutes of data (Gordon, Laumann, Gilmore, et al., 2017; Laumann et al., 2015), the definition of network variants appear to require more data to achieve an adequate reliability. From our estimates, it appears that >70 minutes of (rest or task) data are required to achieve asymptotic reliability for network variants. This added need for data is likely due to the thresholding procedure in defining network variants in either task or rest as spatial correlations maps (Fig. 2A, S1A) achieve high reliability more quickly. Combining task and rest data may be an effective strategy to achieve this amount of data in available datasets with insufficient rest.

In addition to combining rest and task data, our initial evidence suggests that it is possible to either use data from a single task or combine data from multiple tasks to generate network variants. We found no significant differences in the ability to identify network variants across 3 different tasks versus rest, indicating that all of the tasks performed relatively similarly. However, for this analysis we were restricted to using a small amount of data which complicates the interpretation of our analysis. It may be possible that with larger amounts of data or with different tasks, task-specific effects for network variants could be observed.

A final caveat of our analyses is that it is important to match data in terms of both quantity and quality when comparing the reliability of fMRI data. Because reliability of a scan steadily increases with the amount of low-motion data (Elliott et al., 2019; Gordon, Laumann, Gilmore, et al., 2017; Laumann et al., 2015; Noble et al., 2017), it is necessary to match these attributes in order to make accurate comparisons. Factors such as head motion, sampling across sessions, or processing may systematically influence estimates of functional connectivity. In the current study, we found similar levels of motion between task (Mean FD = .129) and rest (Mean FD = .133), and as described in the methods data was approximately sampled across sessions. Recent work also highlights the importance of accounting for the presence of task activations when using task data for functional connectivity (Cole et al., 2019). We also found evidence in line with this conclusion, demonstrating that the presence of task activations makes network variants less similar between task and rest states. Therefore, it is important to account for these task activations to improve similarity between task and rest variants (although overall within-state reliability was similar). Thus, it is critical to control for these factors when comparing data across states, participants, and studies.

Lastly, our results also suggest that using a percent threshold to define variants may work better than an absolute cutoff. We found more stability in variants across states defined with a percentile rather than an absolute cutoff. This may be an important consideration to maximize the reliability of variants across individuals.

## 5 Limitations

We also note several limitations to the current study. First, although the MSC dataset has a large amount of resting state data, it is somewhat limited in the amount of available task data, especially for single tasks. This is particularly true of the motor task, which had a high number of frames lost due to movement. Therefore, we only draw limited inferences about task-specific effects on network variants from the current study.

Second, although we found some state-dependent changes in network variants, it was unclear what drove these differences. We did not find systematic patterns associated with particular networks or particular tasks. It is possible that future work with a broader set or more targeted tasks may help to identify these and shed light on state-dependent mechanisms.

Third, several of the analysis decisions in this paper are based on thresholding data which has the potential to systematically change patterns in the data. One example of this is the thresholding of network variants in individuals as the lowest 5% of correlations with group average networks. This decision was based on examining histograms of group correlations and finding the approximate tail of the distribution. In addition, we also demonstrated that the reported results replicate at other thresholds (2.5%, r < .3) showing their robustness to this choice. Another thresholding choice was in the assignment of variants to networks, which used a winner-take-all network assignment procedure based on the spatial overlap of the top 5% of correlations between a seed region and group average network. It should be noted that the network assignments mirrored past work (Seitzman et al., 2019), with higher-level association networks making up the majority of network variants and seedmap-based comparisons (free of thresholding or winner-take-all selection) were highly stable across states (*Fig. S10*). To alleviate these concerns, future studies may consider employing newer techniques such as bagging (Nikolaidis et al., 2020) to improve reliability of thresholded data or “soft” multi-network assignments (Cui et al., 2020) for networks with an ambiguous pattern of connectivity.

Fourth, we defined network variant “units” based on their contiguity. However, it is possible that contiguous variant units contain adjacent, but not homogenous deviant tissue – that is, they may not index a single underlying brain area. Moreover, some variant units may index shifts in the borders of nearby systems, while others may index more drastic functional alterations in an entire isolated region. We conducted analyses at the vertex-level (see *Figures S11, S13*) to address potential inhomogeneities in network variant units. In future work, we hope to expand our understanding of homogenous variant units and their properties.

Fifth is whether these stable trait-like characteristics are linked to mis-localizations in anatomical alignment. In previous work, we have found that areas with gross anatomical deformations during registration are not associated with network variants (Seitzman et al., 2019). However, it is possible that network variants occur in part due to fine-scale anatomical variations (e.g. cell populations) in given regions of the cortex. These anatomical variations would then contribute to functional variability across individuals. Additional investigations of network variants using different methodologies will be needed to further examine their underlying neurobiology.

## 6 Conclusion

Overall, these results suggest that network variants show trait-like stability between multiple states. Network variants measured with task data were reliable (given sufficient data) and appeared in similar locations with similar network assignment to that seen with rest data. There were also more minor state-dependent differences associated with network variants. Jointly this work suggests that combining rest and task data may be a reasonable strategy to identify network variants from datasets with insufficient rest.

## Acknowledgements

We thank Evan M. Gordon for his assistance with materials used in this project. This work was supported by NIH grant R01MH118370 (to C.G.) and NIH grant T32NS047987 (to B.T.K.).

## Supplementary Results

**Supplementary Table 1 (S1).**
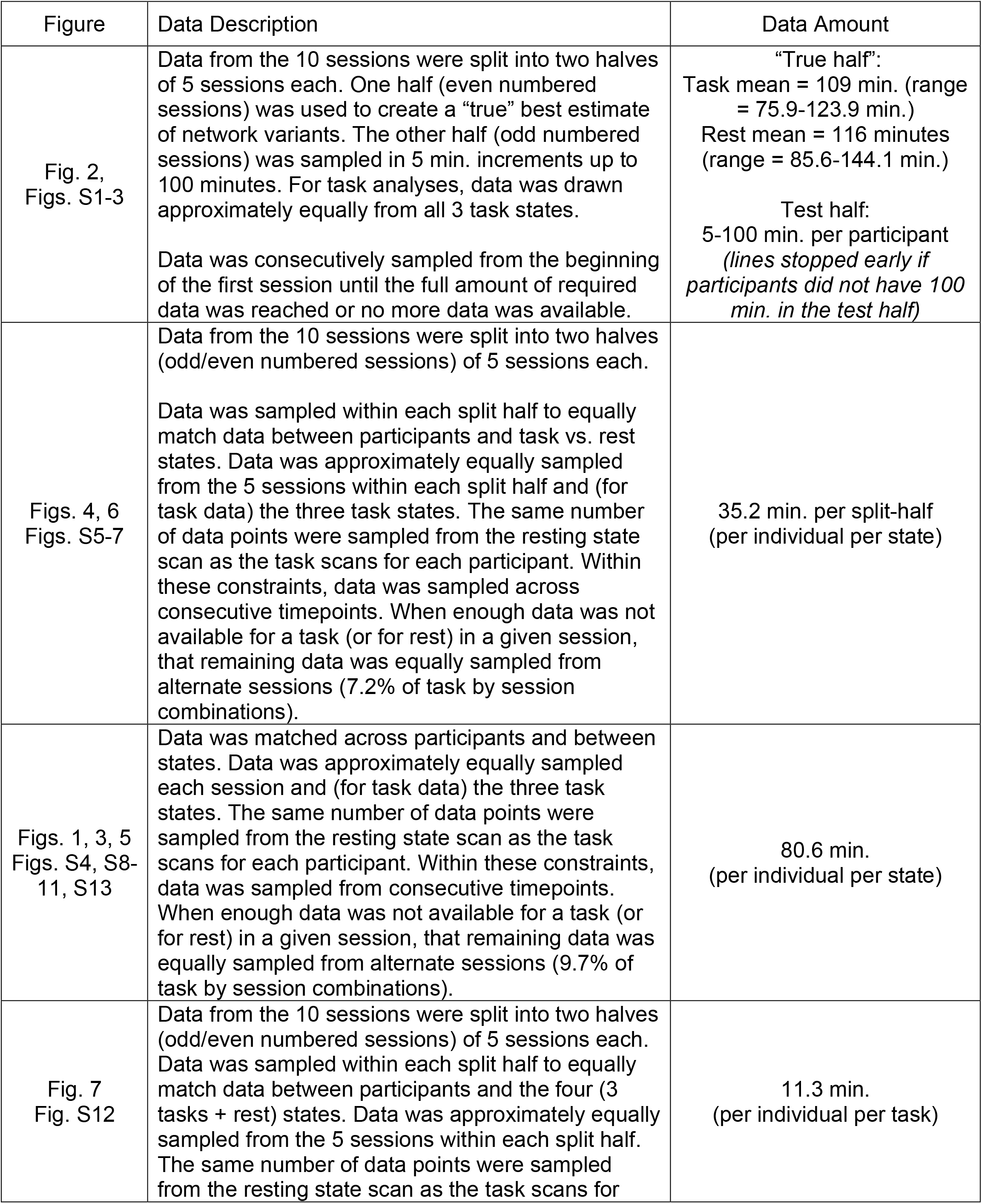
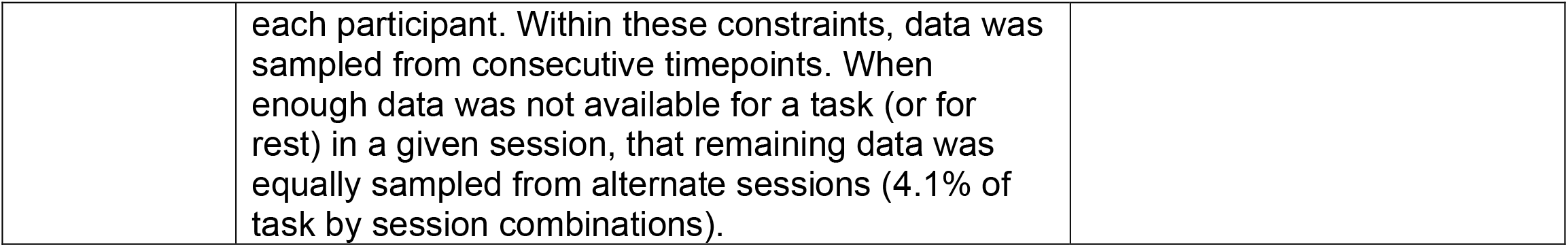
The data sampling procedure for the analysis underlying each figure is outlined.

| Figure | Data Description | Data Amount |
| --- | --- | --- |
| Fig. 2,<br>Figs. S1-3 | Data from the 10 sessions were split into two halves of 5 sessions each. One half (even numbered sessions) was used to create a “true” best estimate of network variants. The other half (odd numbered sessions) was sampled in 5 min. increments up to 100 minutes. For task analyses, data was drawn approximately equally from all 3 task states.<br>Data was consecutively sampled from the beginning of the first session until the full amount of required data was reached or no more data was available. | “True half”:<br>Task mean = 109 min. (range = 75.9-123.9 min.)<br>Rest mean = 116 minutes (range = 85.6-144.1 min.)<br>Test half:<br>5-100 min. per participant<br>*(lines stopped early if participants did not have 100 min. in the test half)* |
| Figs. 4, 6<br>Figs. S5-7 | Data from the 10 sessions were split into two halves (odd/even numbered sessions) of 5 sessions each.<br>Data was sampled within each split half to equally match data between participants and task vs. rest states. Data was approximately equally sampled from the 5 sessions within each split half and (for task data) the three task states. The same number of data points were sampled from the resting state scan as the task scans for each participant. Within these constraints, data was sampled across consecutive timepoints. When enough data was not available for a task (or for rest) in a given session, that remaining data was equally sampled from alternate sessions (7.2% of task by session combinations). | 35.2 min. per split-half<br>(per individual per state) |
| Figs. 1, 3, 5<br>Figs. S4, S8-11, S13 | Data was matched across participants and between states. Data was approximately equally sampled each session and (for task data) the three task states. The same number of data points were sampled from the resting state scan as the task scans for each participant. Within these constraints, data was sampled from consecutive timepoints. When enough data was not available for a task (or for rest) in a given session, that remaining data was equally sampled from alternate sessions (9.7% of task by session combinations). | 80.6 min.<br>(per individual per state) |
| Fig. 7<br>Fig. S12 | Data from the 10 sessions were split into two halves (odd/even numbered sessions) of 5 sessions each. Data was sampled within each split half to equally match data between participants and the four (3 tasks + rest) states. Data was approximately equally sampled from the 5 sessions within each split half. The same number of data points were sampled from the resting state scan as the task scans for | 11.3 min.<br>(per individual per task) |
|  | each participant. Within these constraints, data was sampled from consecutive timepoints. When enough data was not available for a task (or for rest) in a given session, that remaining data was equally sampled from alternate sessions (4.1% of task by session combinations). |  |

**Supplementary Table 2 (S2).** The total number of variants is shown for task and rest states for each participant at a 5% threshold. These counts reflect variants that were excluded for being in low signal regions, small size (< 50 vertices), or overlapping more than 50% with the group network template.

| Subject | Rest | Task |
| --- | --- | --- |
| MSC01 | 16 | 17 |
| MSC02 | 13 | 11 |
| MSC03 | 5 | 7 |
| MSC04 | 16 | 14 |
| MSC05 | 12 | 16 |
| MSC06 | 9 | 11 |
| MSC07 | 17 | 12 |
| MSC09 | 12 | 14 |
| MSC10 | 14 | 10 |
| Total | 114 | 112 |

**Supplementary Table 3 (S3).** The total number of variant vertices is shown for task and rest states for each participant at a r < .3 threshold. These counts reflect variant vertices that were excluded for being in low signal regions and small size (< 50 vertices). There were no significant differences in the number of vertices between states *t*(8) = .044, *p* = .9661, *d* =.01.

| Subject | Rest | Task |
| --- | --- | --- |
| MSC01 | 2192 | 1975 |
| MSC02 | 1007 | 611 |
| MSC03 | 1050 | 1332 |
| MSC04 | 343 | 938 |
| MSC05 | 954 | 567 |
| MSC06 | 1529 | 1579 |
| MSC07 | 927 | 1128 |
| MSC09 | 475 | 532 |
| MSC10 | 746 | 604 |
| Total | 9233 | 9266 |

**Figure S1.**
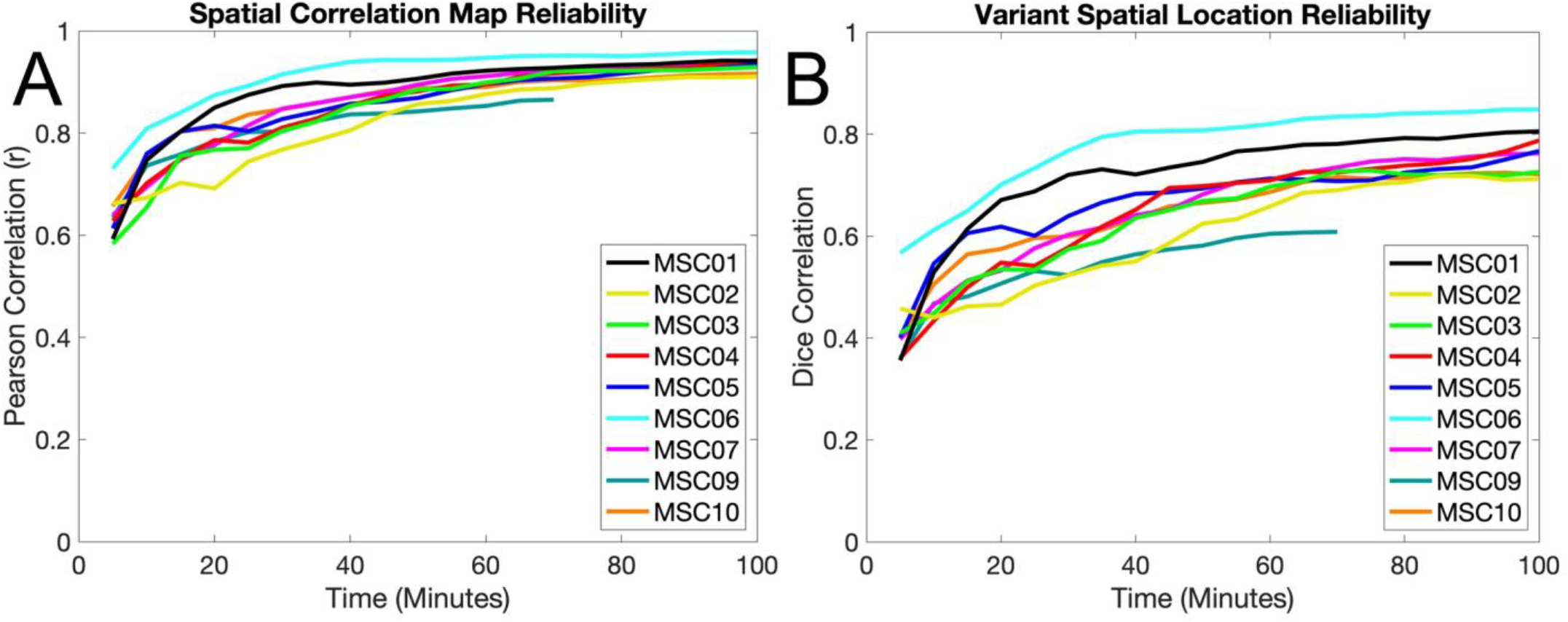
Reliability of network variants in rest data. As for the task data in Figure 2, sessions were split into two halves and compared. All available data was taken from one half and treated as the best estimate of “true” network variants. The other “test” half was sampled in 5 min. increments. (A) Individual-to-group spatial correlation maps were compared using Pearson correlation. (B) Binarized network variant maps (lowest 5% of correlations) were compared using Dice correlation. Note that MSC09 (70 minutes) did not reach 100 minutes of data in their test half and thus has a shorter line than the rest of the participants.

**Figure S2.**
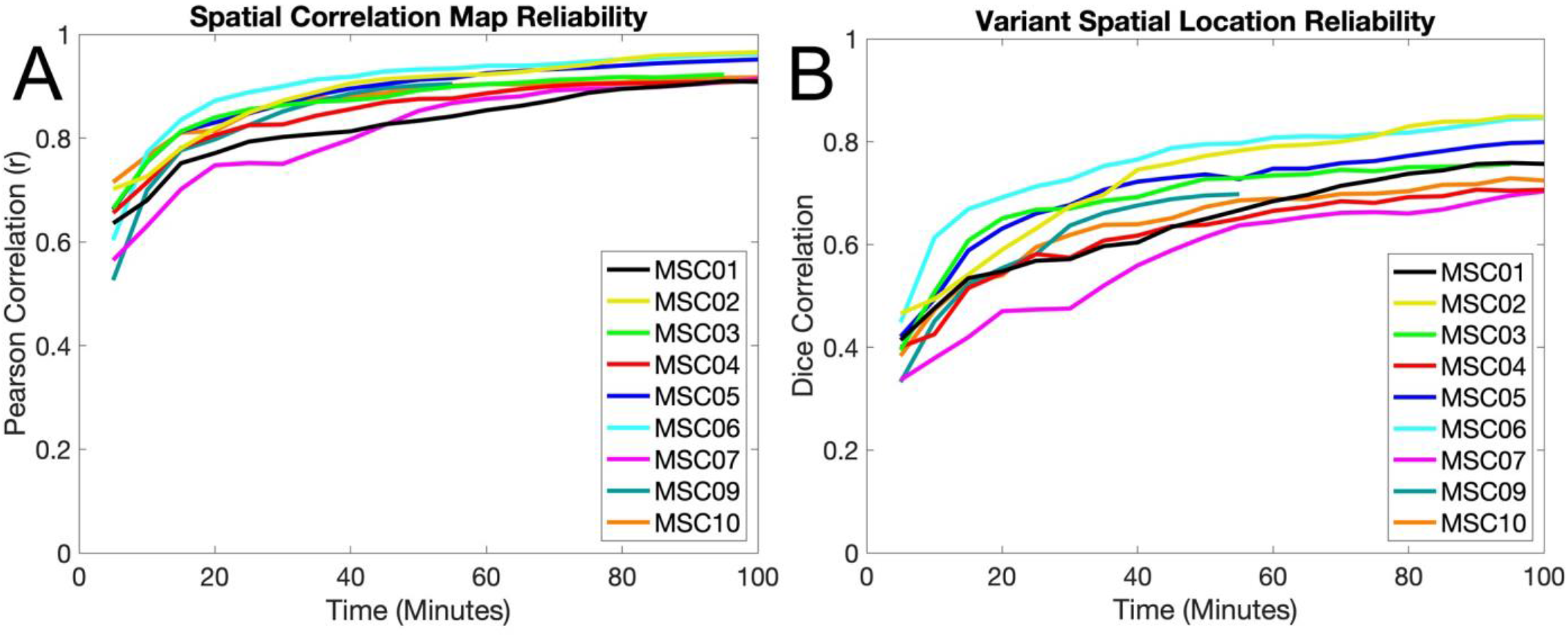
Reliability of non-residualized data (i.e., data without task activations removed), as in Figure 2. Sessions were split into two halves and compared. All available non-residualized data was taken from one half and treated as the best estimate of “true” network variants. The other “test” half of non-residualized data was sampled in 5 min. increments. (A) Individual-to-group spatial correlation maps were compared using Pearson correlation. (B) Binarized network variant maps (lowest 5% of correlations) were compared using Dice correlation. Note that MSC09 (55 minutes) and MSC03 (95 minutes) did not have 100 minutes of data in their test half, leading to shorter lines. Similar reliability was seen for residualized and non-residualized task data.

**Figure S3.**
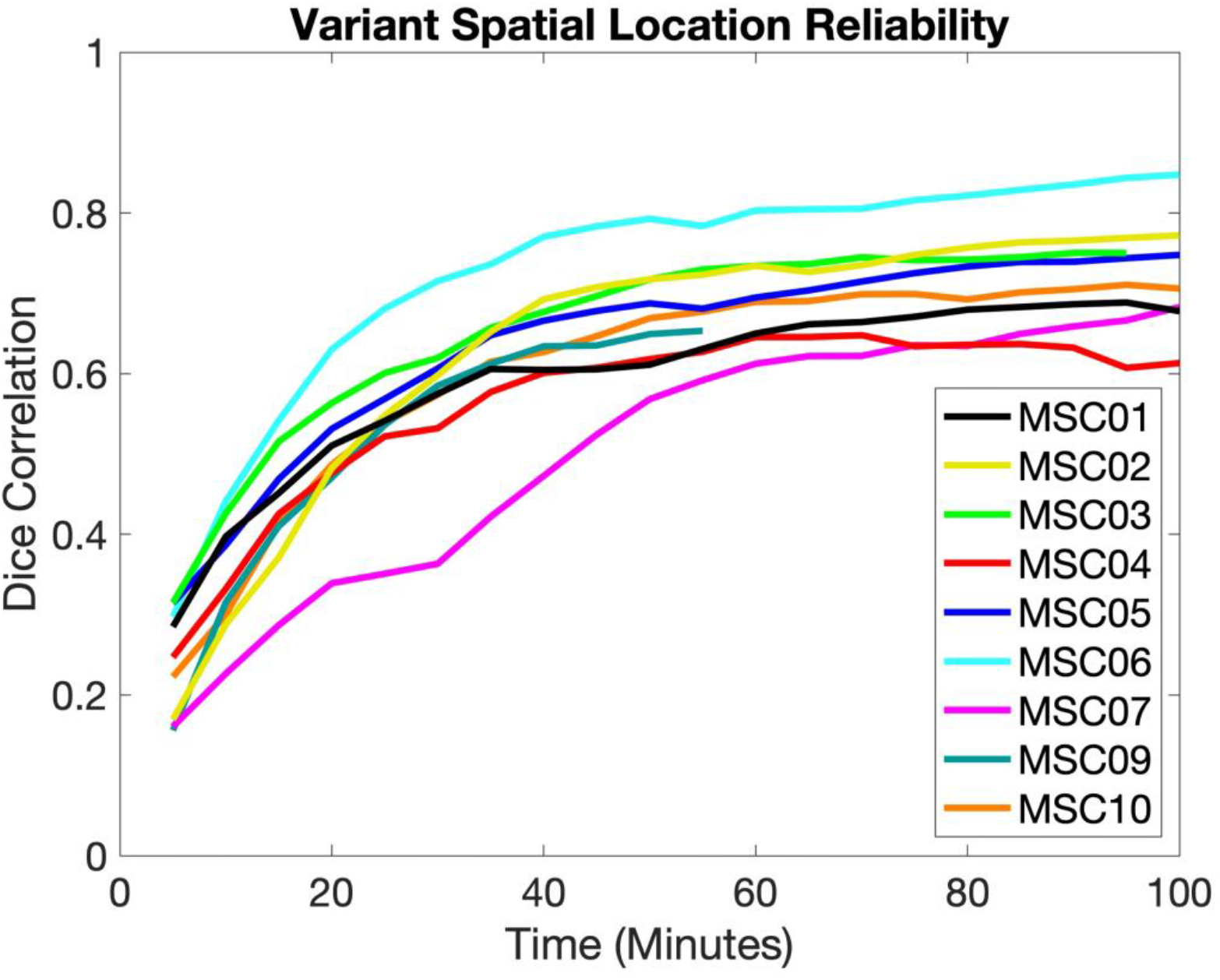
As for the task data in Figure 2, sessions were split into two halves and compared. All available data was taken from one half and treated as the best estimate of “true” network variants. The other “test” half was sampled in 5 min. increments. Binarized network variant maps (r < 0.3 between the individual and group average) were compared using Dice correlation. Note that MSC09 (70 minutes) did not reach 100 minutes of data in their test half and thus have shorter lines than the other participants.

**Figure S4.**
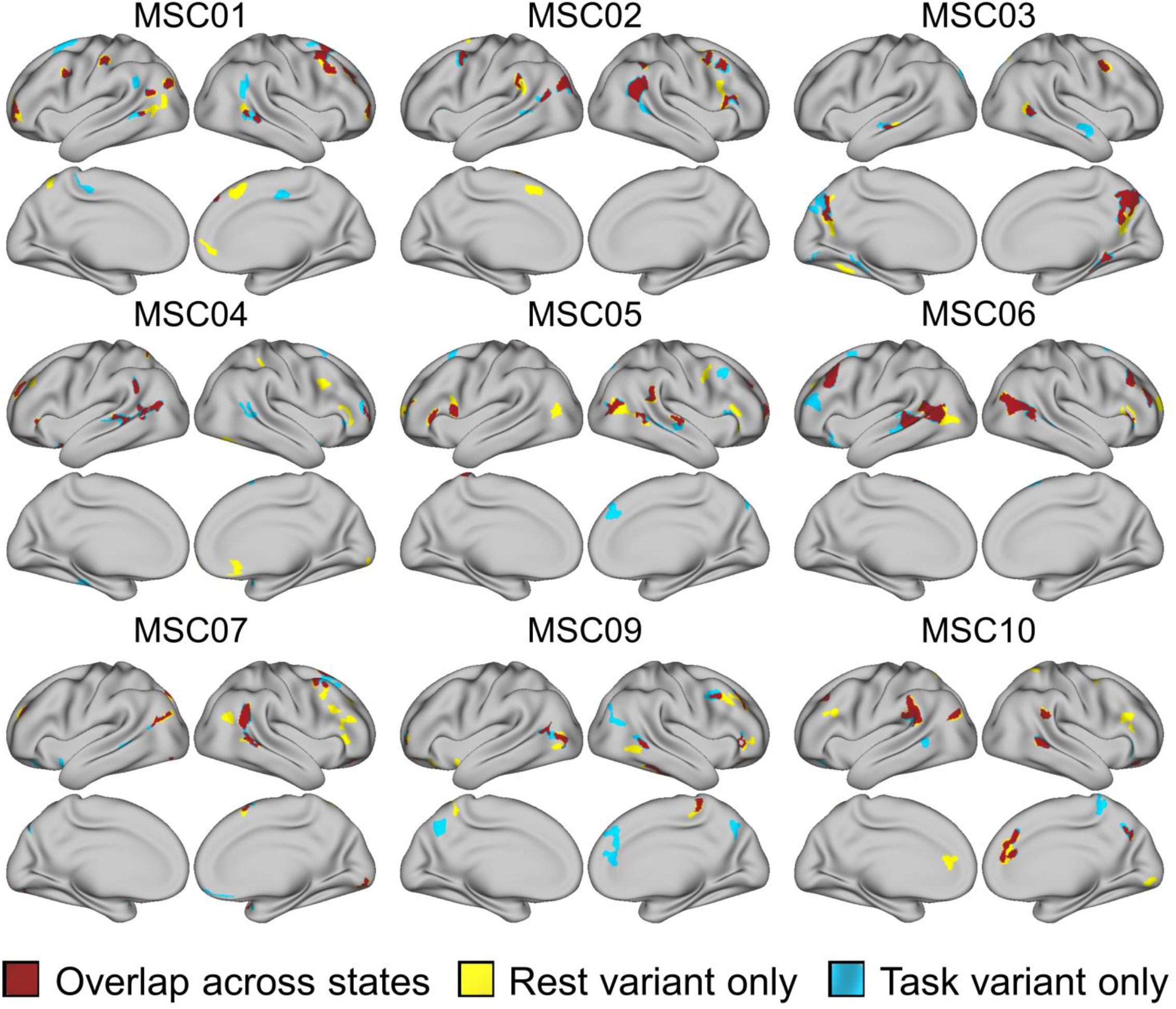
The overlap in variants between task and rest is shown for all included MSC subjects (using a 5% network variant threshold). Areas shaded in yellow represent variants that are only observed during the resting state, areas shaded in blue represent areas only observed in the task state, and areas shaded in red represent areas where variants are present in both task and rest states (overlapping).

**Figure S5.**
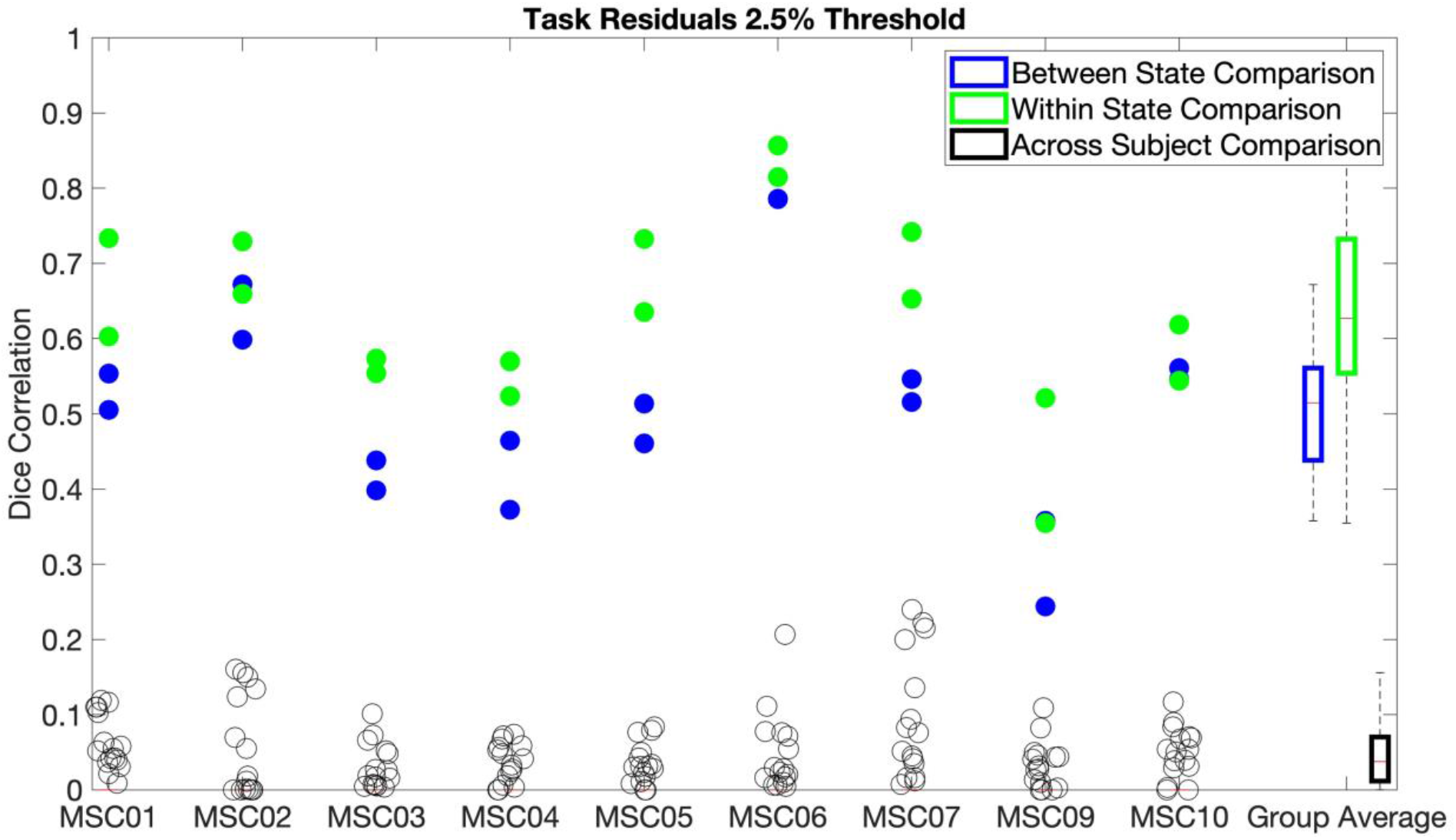
The Dice correlation of the overlap of vertices containing network variants between states and within states is plotted for variants thresholded at the 2.5% lowest values of each individual’s spatial correlation map. The value for each individual participant is plotted for all of the comparisons, and the value of every across participant comparison is also plotted. Two dots are present for the between and within state comparisons as both pairs of split-halves were used (see sections 2.9.4 and 2.9.5). The results at the 2.5% threshold showed that variants were more likely to overlap between states (*M* = .518, *SD* = .139) than across subjects (*M* = .049, *SD* = .02, *t*(8) = 10.56 *p* < .0001, *d* = 3.52). Variants were also were significantly more likely to overlap within states (*M* = .635, *SD* = .115) than between states (*M* = .518, *SD* = .139, *t*(8) = 6.1, *p* = .0003, *d* = 2.03; see Figure 4). As with the 5% threshold, the magnitude of the difference between states was smaller than the magnitude of the difference across subjects.

**Figure S6.**
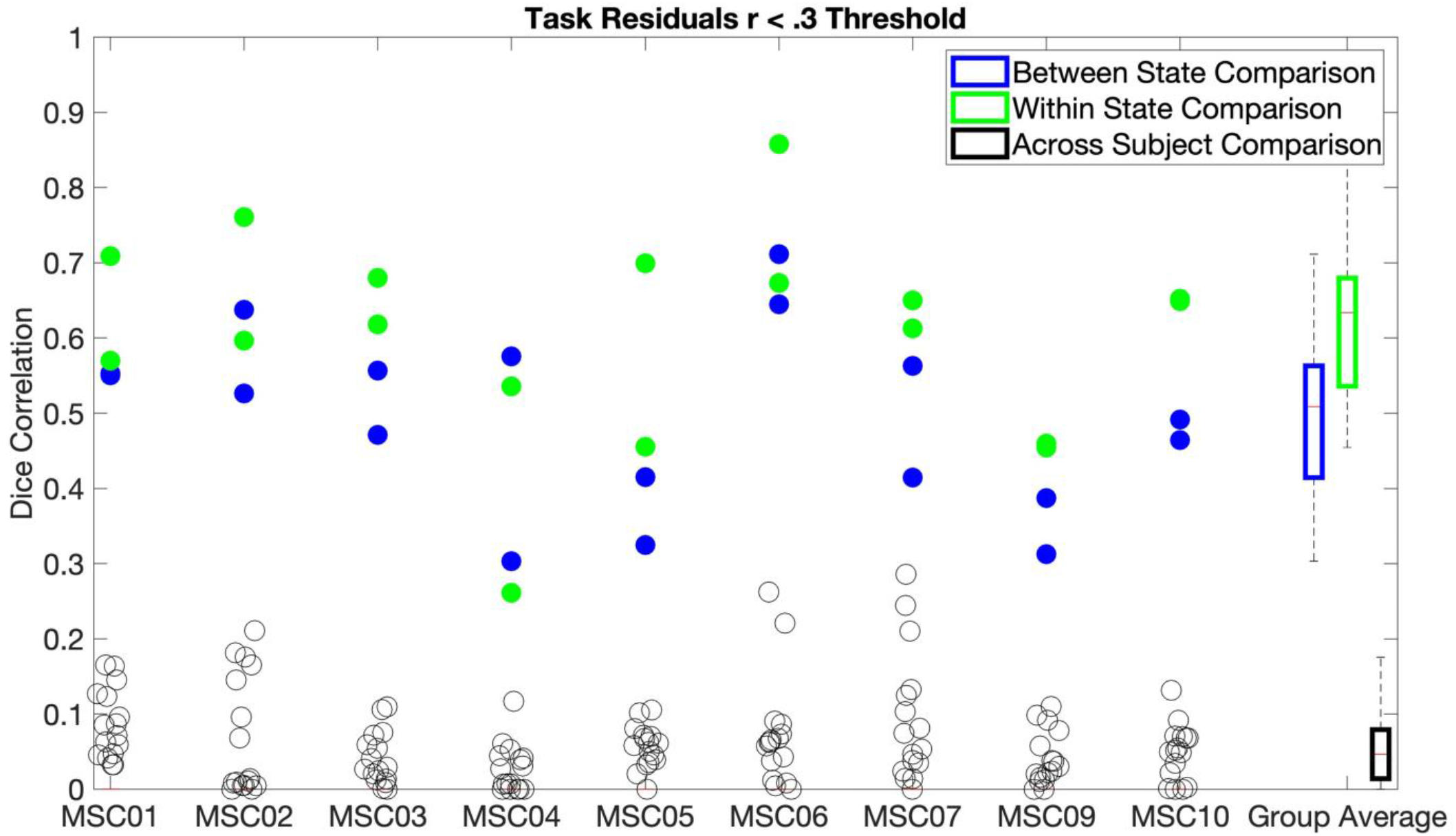
The Dice correlation of the overlap of vertices containing network variants between states and within states is plotted for variants thresholded at values of r < .3 of each individual’s spatial correlation map. The value for each individual participant is plotted for all of the comparisons, and the value of every across participant comparison is also plotted. Two dots are present for the between and within state comparisons as both pairs of split-halves were used (see sections 2.9.4 and 2.9.5). The results at the r < .3 threshold showed that variants were more likely to overlap between states (*M* = .495, *SD* = .103) than across subjects (*M* = .059, *SD* = .022, *t*(8) = 13.94, *p* < .0001, *d* = 4.65). Variants were also were significantly more likely to overlap within (*M* = .605, *SD* = .113) than between states (*M* = .495, *SD* = .103, *t*(8) = 4.75, *p* = .001, *d* = 1.58; see Figure 4). As with the 5% threshold, the magnitude of the difference between states was smaller than the magnitude of the difference across subjects.

**Figure S7.**
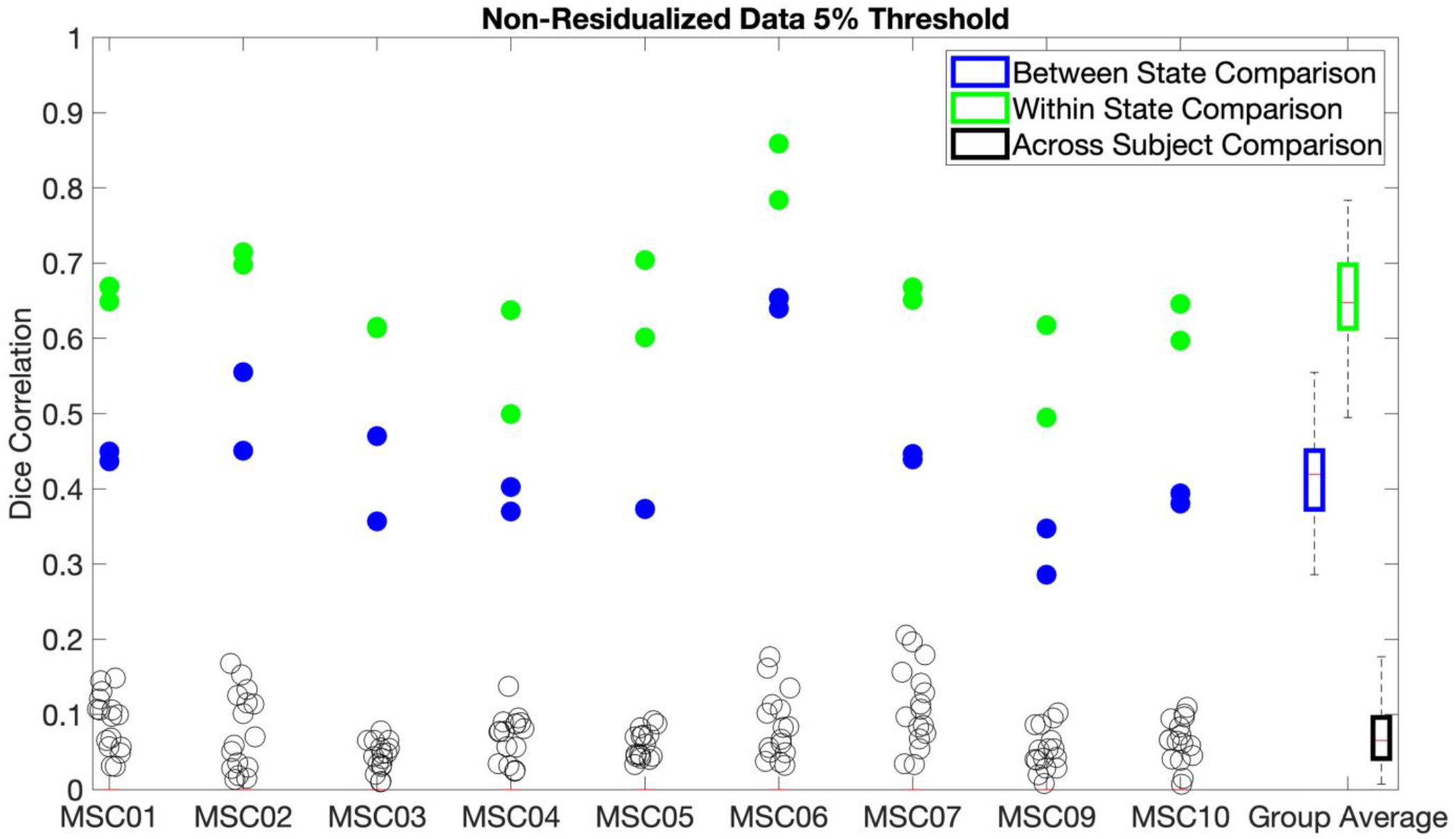
The Dice correlation of the overlap of vertices containing network variants in non-residualized task data between states and within states is plotted for variants thresholded at the 5% lowest values of each individual’s spatial correlation map. The value for each individual participant is plotted for all of the comparisons, and the value of every across participant comparison is also plotted. Two dots are present for the between and within state comparisons as both pairs of split-halves were used (see sections 2.9.4 and 2.9.5). As with the findings from the main manuscript, results for the non-residualized data showed that variants were more likely to overlap between states (*M* = .435, *SD* = .095) than across subjects (*M* = .072, *SD* = .02, *t*(8) = 12.46, *p* < .0001, *d* = 4.15). Variants were also were significantly more likely to overlap within states (*M* = .651, *SD* = .079) than between states (*M* = .435, *SD* = .095, *t*(8) = 20.31 *p* < .0001, *d* = 6.77; see Figure 4). However, direct comparisons showed that non-residualized data had significantly lower between-state stability than residualized data (*p* < .0001, *d* = 2.7). Thus, it appears that removing task activations via regression provides more cross-state stability than when task activations are left in the data.

**Figure S8.**
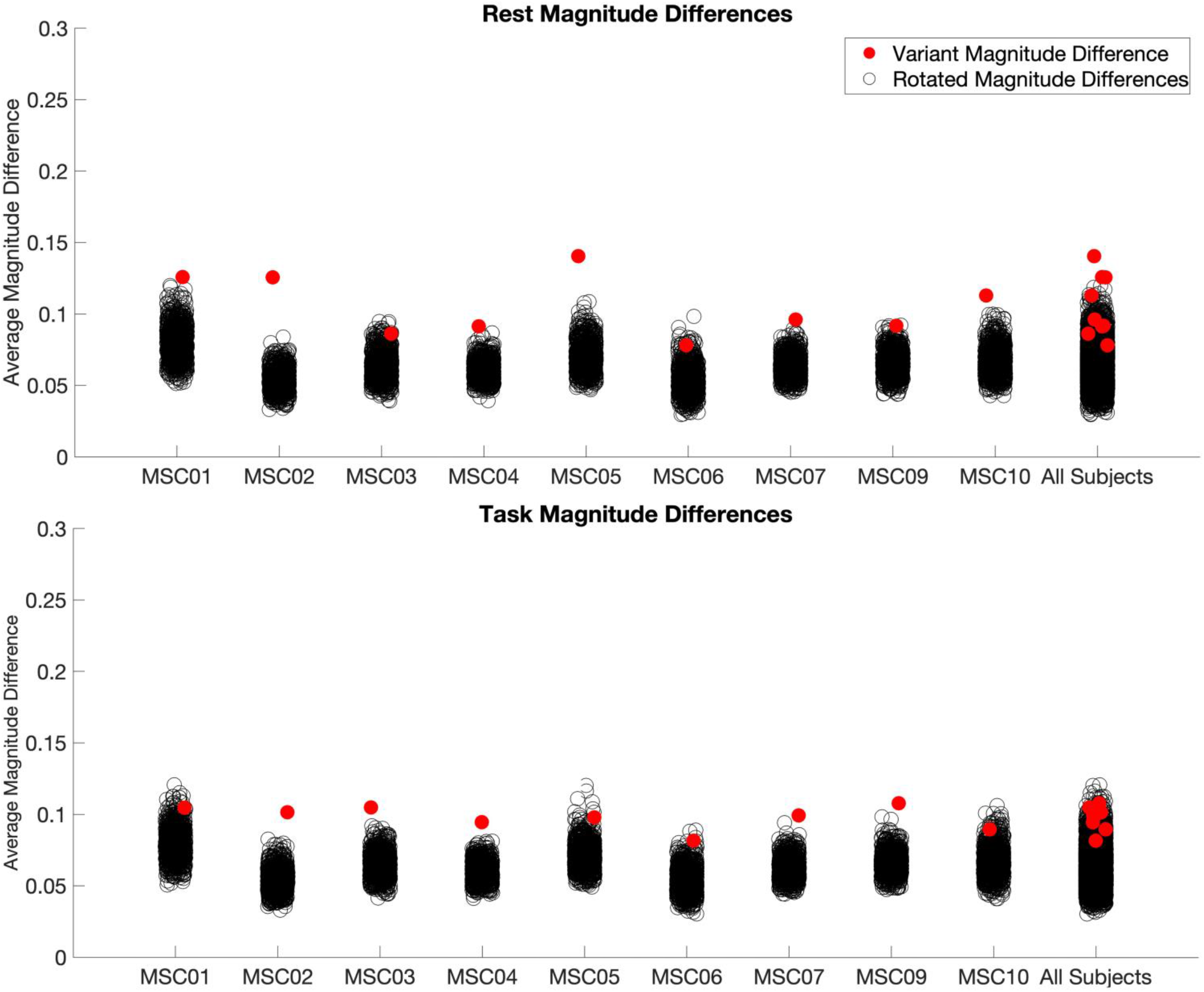
The difference in magnitude of correlations with the group average networks between variants in rest and task states. In this analysis, the spatial correlations of the vertices identified as network variants during one state were subtracted from the spatial correlations of the same locations during the alternate state. Then the mean of the absolute value of this difference was calculated to obtain a mean difference in magnitude of the correlations between states (see Figure 3C). The magnitude difference is shown separately for variants identified in rest (top) and variants identified in task (bottom). To determine whether these values were different from what would be expected by chance (i.e., relative to other areas of the cortex), the network variants for each participant were rotated 1000 times within each hemisphere. The same operation was then performed for each rotation. Red dots represent the average (absolute) magnitude difference of the observed network variants and the black dots represent the average (absolute) magnitude difference of randomly rotated variants. Network variants showed more variation than would be expected by chance for most participants. As can be seen in Figure 3C, most locations show small deviations (< 0.1), but some locations show larger differences, likely driving this effect.

**Figure S9.**
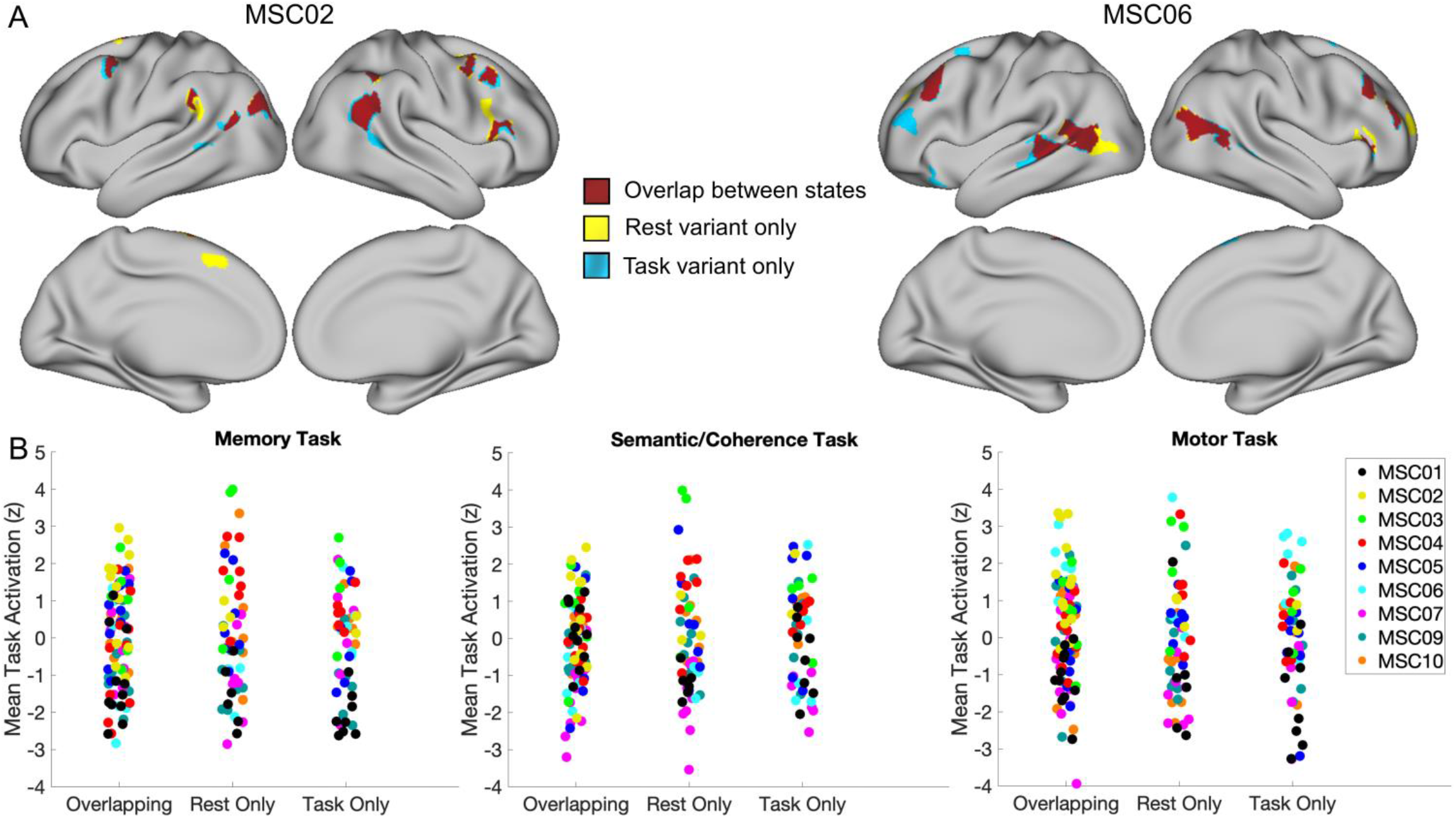
Contribution of task activations to state-dependence in network variants. Variant sub-units that either occur in both states (overlapping; red), only in rest (yellow), or only in task (blue) are shown on the cortical surface for MSC02 and MSC06 (A). For each of these sub-units, the mean task activation was calculated for each variant in each task (B). There were no significant differences in mean task activations across the different sub-units in each task (*p* < .05).

**Figure S10.**
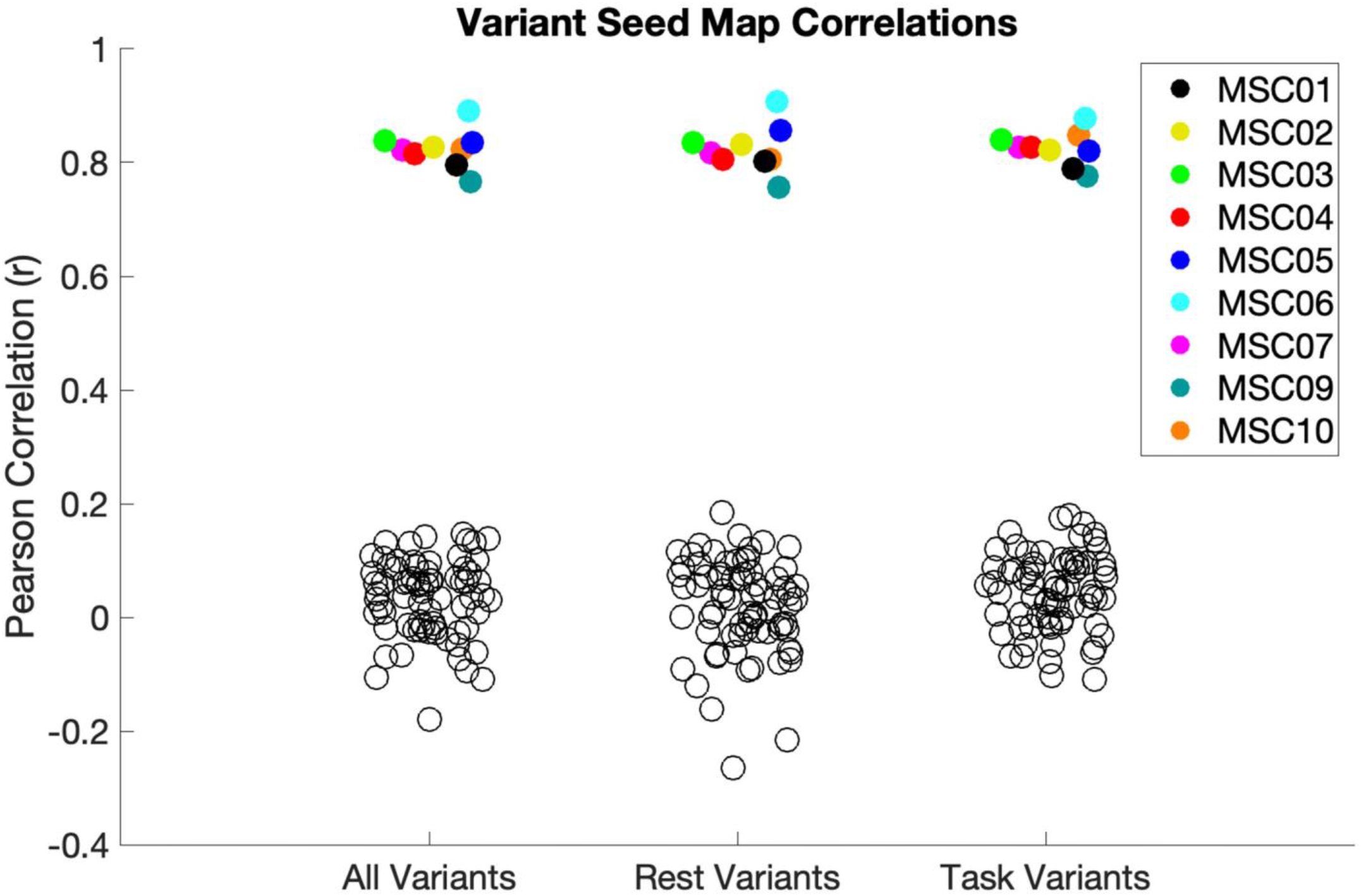
Comparison of seedmaps for network variants. The average spatial correlations for the variant seed maps between task and rest states is plotted (average per subject). Plots are shown for all variants and for variant estimates made separately during task or rest. Empty black circles show comparisons between network variant seedmaps and the same locations in other subjects. Variant seedmaps were quite similar between states (*M* = .824) and much higher than seen across subjects (*M* = .034), *t*(8) = 33.07, *p* < .0001, *d* = 11.02.

**Figure S11.**
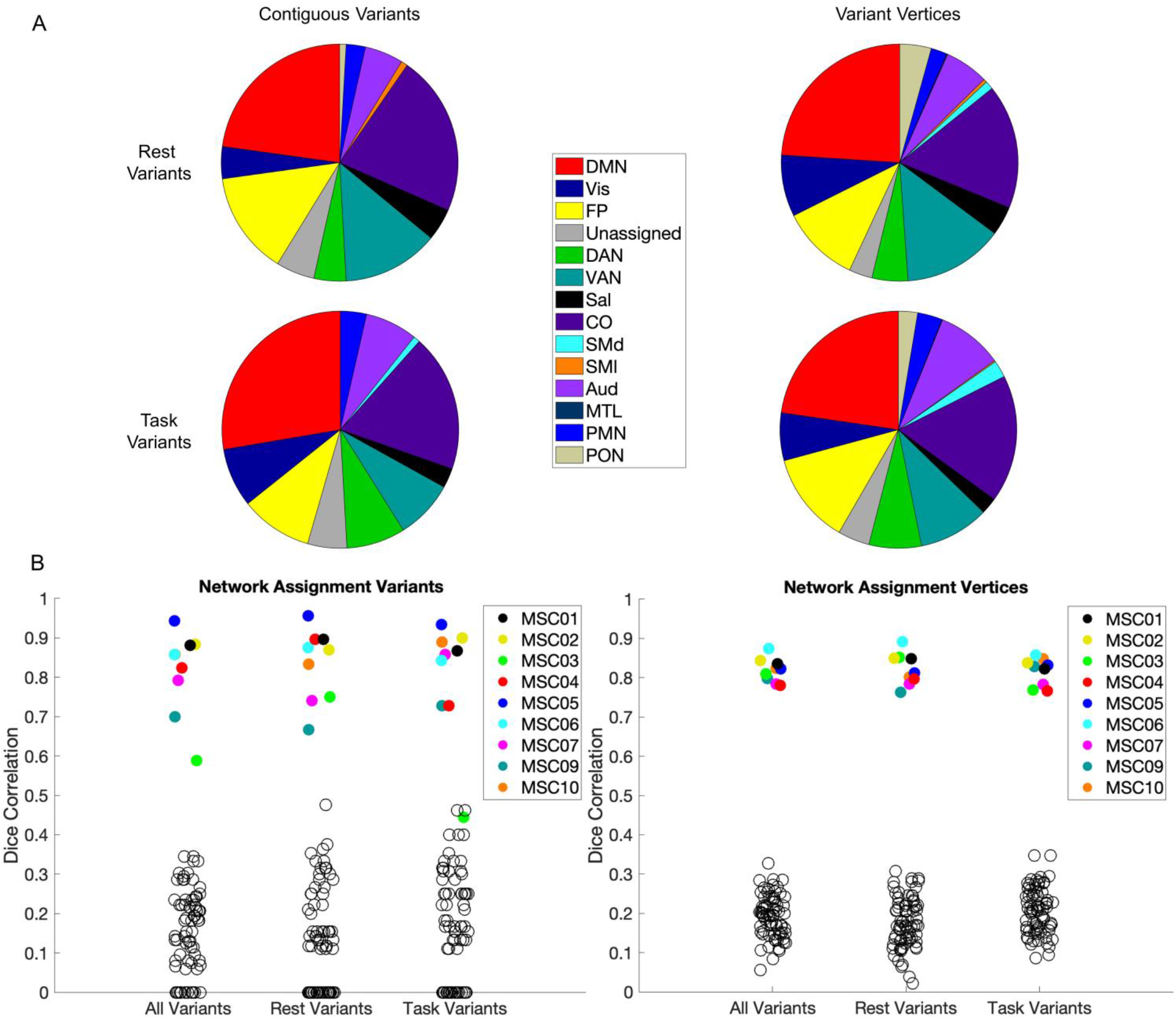
Comparison of network assignments conducted for contiguous variant units (left) or for each vertex in a variant separately (right). (A) Distribution of network variant assignments during task and rest states. (B) The likelihood network variants are assigned to the same network across states (shown for variant locations identifed based on both task and rest states as well as broken up by individual states). Empty black circles represent the likelihood of variant locations assigning to the same network in other subjects. As with contiguous variants, vertices were much more likely to assign to the same network between states (*M* = .819, *SD* = .03) than across subjects (*M* = .193, *SD* = .028), *t*(8) = 41.69, *p* <. 0001, *d* = 13.9.

**Figure S12.**
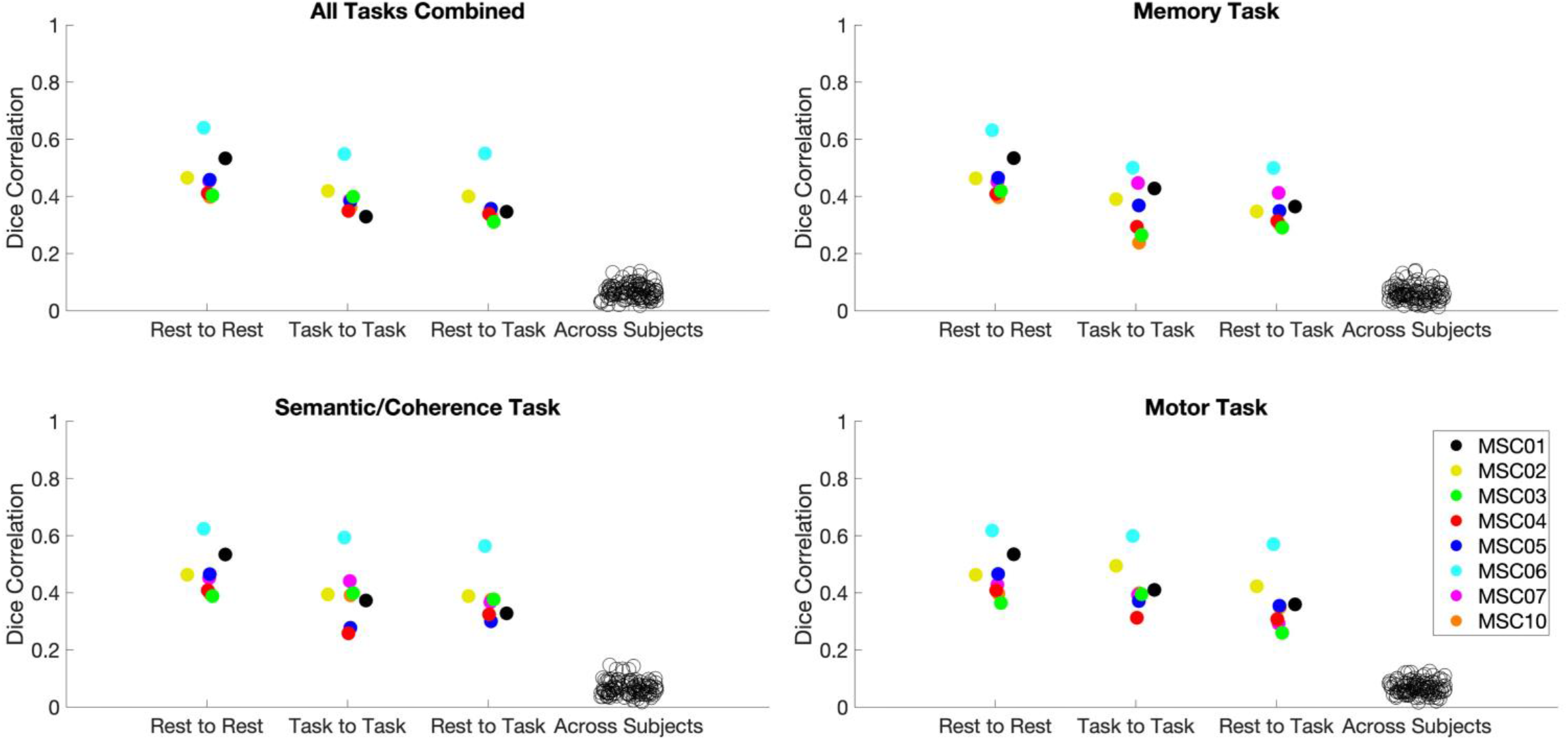
Comparisons for network variant stability across individual tasks at a 2.5% threshold. Dice correlations for the stability of all 4 tasks are shown using a 2.5% threshold of the lowest correlations with the group average to define network variants. The pattern shown is similar to that observed in Figure 7. As in section 3.5, a within-subject ANOVA was used to test for significant differences in the stability of variants between tasks using a 2.5% threshold for defining variants. A model with one independent variable (Task) was specified for the rest to task comparisons across all 4 tasks. This model was not significant, *F*(3,21) = 1.193, *p* = .336, *d* = .83, indicating that there were no differences in stability between states in any of the four tasks. These results are the same as those observed at the 5% threshold, providing insufficient evidence that there are task-specific effects on the stability of network variants between states.

**Figure S13.**
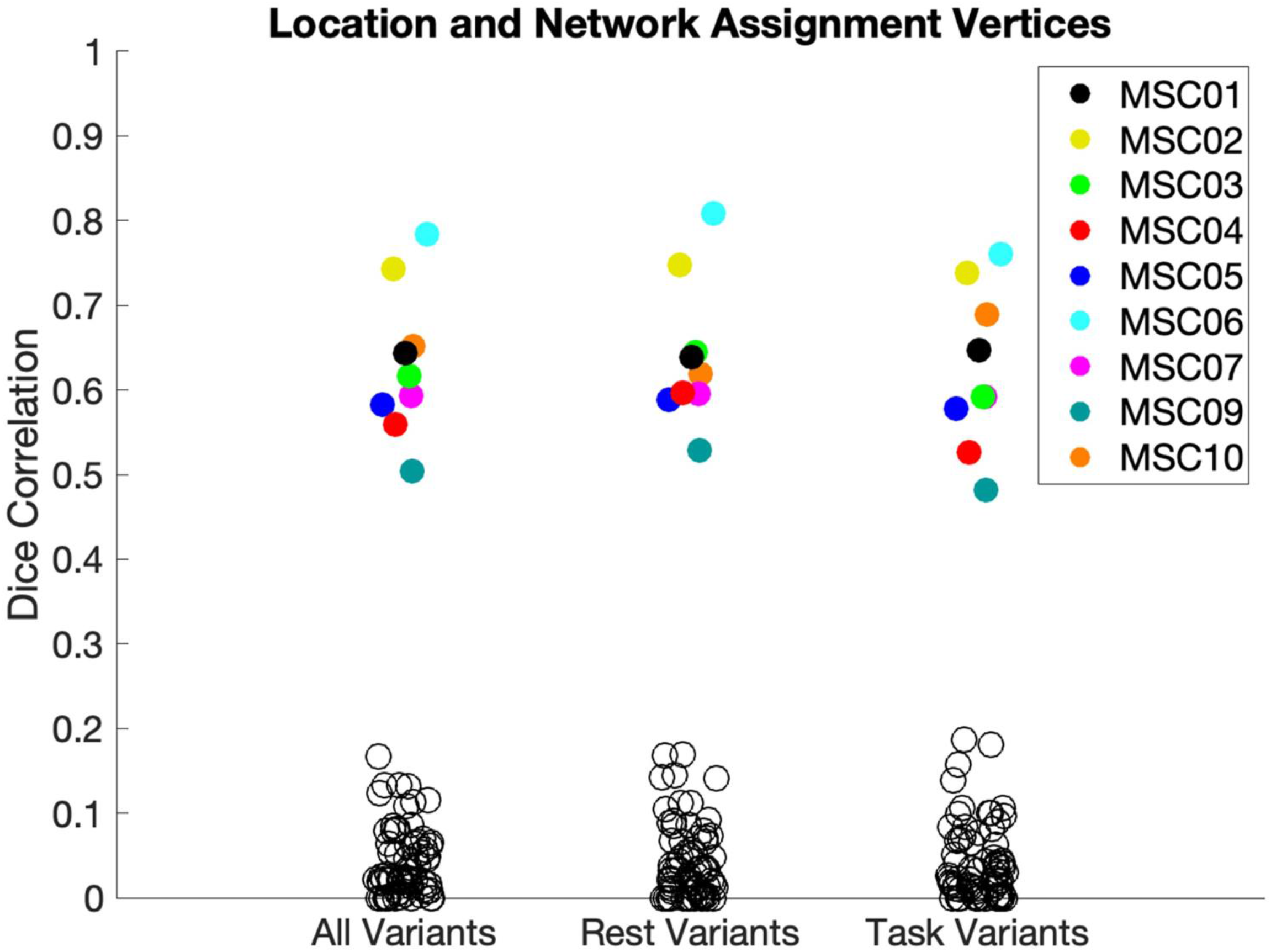
Dice correlations showing the likelihood of the vertices composing each network variant both spatially overlapping (see Figure 4) and assigning to the same network (see Figure 5). The empty black circles represent the likelihood of these vertices spatially overlapping and assigning to the same network in the opposite state across all subjects. Variant vertices which overlapped with the consensus networks were excluded from analysis and vertices with an assignment lower than the threshold reported in section 2.12.1 were assigned to an “unknown” network. Variants were much more likely to spatially overlap and assign to the same network between states (*M* =.631) than across subjects (*M* =.041), *t*(8) = 20.33, *p* < .0001, *d* = 6.78. Note that this number is higher than the number reported for spatial overlap alone between states in Figure 4 because this analysis used the same amount of data per subject as used in Figure 5 (80.6 minutes) versus the amount used in Figure 4 (35.2 minutes) which increased the mean spatial overlap between states to .769.

## References

Al-Aidroos, N., Said, C. P., & Turk-Browne, N. B. (2012). Top-down attention switches coupling between low-level and high-level areas of human visual cortex. Proceedings of the National Academy of Sciences, 109(36), 14675–14680.

Barch, D. M., Burgess, G. C., Harms, M. P., Petersen, S. E., Schlaggar, B. L., Corbetta, M., Glasser, M. F., Curtiss, S., Dixit, S., & Feldt, C. (2013). Function in the human connectome: Task-fMRI and individual differences in behavior. Neuroimage, 80, 169–189.

Biswal, B., Zerrin Yetkin, F., Haughton, V. M., & Hyde, J. S. (1995). Functional connectivity in the motor cortex of resting human brain using echo-planar MRI. Magnetic Resonance in Medicine, 34(4), 537– 541.

Braga, R. M., & Buckner, R. L. (2017). Parallel interdigitated distributed networks within the individual estimated by intrinsic functional connectivity. Neuron, 95(2), 457–471.

Braga, R. M., DiNicola, L. M., Becker, H. C., & Buckner, R. L. (2019). Situating the left-lateralized language network in the broader organization of multiple specialized large-scale distributed networks. Journal of Neurophysiology.

Cho, J. W., Korchmaros, A., Vogelstein, J. T., Milham, M., & Xu, T. (2020). Impact of Concatenating fMRI Data on Reliability for Functional Connectomics. BioRxiv.

Cole, M. W., Bassett, D. S., Power, J. D., Braver, T. S., & Petersen, S. E. (2014). Intrinsic and task-evoked network architectures of the human brain. Neuron, 83(1), 238–251.

Cole, M. W., Ito, T., Schultz, D., Mill, R., Chen, R., & Cocuzza, C. (2019). Task activations produce spurious but systematic inflation of task functional connectivity estimates. NeuroImage, 189, 1–18.

Cui, Z., Li, H., Xia, C. H., Larsen, B., Adebimpe, A., Baum, G. L., Cieslak, M., Gur, R. E., Gur, R. C., & Moore, T. M. (2020). Individual variation in functional topography of association networks in youth. Neuron.

Dale, A. M., Fischl, B., & Sereno, M. I. (1999). Cortical surface-based analysis: I. Segmentation and surface reconstruction. Neuroimage, 9(2), 179–194.

Dice, L. R. (1945). Measures of the amount of ecologic association between species. Ecology, 26(3), 297–302.

DiNicola, L. M., Braga, R. M., & Buckner, R. L. (2020). Parallel distributed networks dissociate episodic and social functions within the individual. Journal of Neurophysiology, 123(3), 1144–1179.

Dosenbach, N. U., Fair, D. A., Miezin, F. M., Cohen, A. L., Wenger, K. K., Dosenbach, R. A., Fox, M. D., Snyder, A. Z., Vincent, J. L., & Raichle, M. E. (2007). Distinct brain networks for adaptive and stable task control in humans. Proceedings of the National Academy of Sciences, 104(26), 11073–11078.

Elliott, M. L., Knodt, A. R., Cooke, M., Kim, M. J., Melzer, T. R., Keenan, R., Ireland, D., Ramrakha, S., Poulton, R., & Caspi, A. (2019). General Functional Connectivity: Shared features of resting-state and task fMRI drive reliable and heritable individual differences in functional brain networks. NeuroImage, 189, 516–532.

Fair, D. A., Miranda-Dominguez, O., Snyder, A. Z., Perrone, A., Earl, E. A., Van, A. N., Koller, J. M., Feczko, E., Tisdall, M. D., & van der Kouwe, A. (2020). Correction of respiratory artifacts in MRI head motion estimates. NeuroImage, 208, 116400.

Fair, D. A., Schlaggar, B. L., Cohen, A. L., Miezin, F. M., Dosenbach, N. U., Wenger, K. K., Fox, M. D., Snyder, A. Z., Raichle, M. E., & Petersen, S. E. (2007). A method for using blocked and event-related fMRI data to study “resting state” functional connectivity. Neuroimage, 35(1), 396–405.

Finn, E. S., & Bandettini, P. A. (2020). Movie-watching outperforms rest for functional connectivity-based prediction of behavior. BioRxiv.

Finn, E. S., Shen, X., Scheinost, D., Rosenberg, M. D., Huang, J., Chun, M. M., Papademetris, X., & Constable, R. T. (2015). Functional connectome fingerprinting: Identifying individuals using patterns of brain connectivity. Nature Neuroscience, 18(11), 1664.

Fischl, B., Salat, D. H., Busa, E., Albert, M., Dieterich, M., Haselgrove, C., Van Der Kouwe, A., Killiany, R., Kennedy, D., & Klaveness, S. (2002). Whole brain segmentation: Automated labeling of neuroanatomical structures in the human brain. Neuron, 33(3), 341–355.

Glass, L. (1969). Moire effect from random dots. Nature, 223(5206), 578.

Glasser, M. F., Sotiropoulos, S. N., Wilson, J. A., Coalson, T. S., Fischl, B., Andersson, J. L., Xu, J., Jbabdi, S., Webster, M., & Polimeni, J. R. (2013). The minimal preprocessing pipelines for the Human Connectome Project. Neuroimage, 80, 105–124.

Gordon, E. M., Laumann, T. O., Adeyemo, B., Gilmore, A. W., Nelson, S. M., Dosenbach, N. U., & Petersen, S. E. (2017). Individual-specific features of brain systems identified with resting state functional correlations. NeuroImage, 146, 918–939.

Gordon, E. M., Laumann, T. O., Adeyemo, B., Huckins, J. F., Kelley, W. M., & Petersen, S. E. (2016). Generation and evaluation of a cortical area parcellation from resting-state correlations. Cerebral Cortex, 26(1), 288–303.

Gordon, E. M., Laumann, T. O., Adeyemo, B., & Petersen, S. E. (2017). Individual variability of the system-level organization of the human brain. Cerebral Cortex, 27(1), 386–399.

Gordon, E. M., Laumann, T. O., Gilmore, A. W., Newbold, D. J., Greene, D. J., Berg, J. J., Ortega, M., Hoyt-Drazen, C., Gratton, C., & Sun, H. (2017). Precision functional mapping of individual human brains. Neuron, 95(4), 791–807.

Gratton, C., Dworetsky, A., Coalson, R. S., Adeyemo, B., Laumann, T. O., Wig, G. S., Kong, T. S., Gratton, G., Fabiani, M., & Barch, D. M. (2020). Removal of high frequency contamination from motion estimates in single-band fMRI saves data without biasing functional connectivity. NeuroImage, 116866.

Gratton, C., Kraus, B. T., Greene, D. J., Gordon, E. M., Laumann, T. O., Nelson, S. M., Dosenbach, N. U., & Petersen, S. E. (2019). Defining Individual-Specific Functional Neuroanatomy for Precision Psychiatry. Biological Psychiatry.

Gratton, C., Laumann, T. O., Gordon, E. M., Adeyemo, B., & Petersen, S. E. (2016). Evidence for two independent factors that modify brain networks to meet task goals. Cell Reports, 17(5), 1276–1288.

Gratton, C., Laumann, T. O., Nielsen, A. N., Greene, D. J., Gordon, E. M., Gilmore, A. W., Nelson, S. M., Coalson, R. S., Snyder, A. Z., & Schlaggar, B. L. (2018). Functional brain networks are dominated by stable group and individual factors, not cognitive or daily variation. Neuron, 98(2), 439–452.

Greene, D. J., Koller, J. M., Hampton, J. M., Wesevich, V., Van, A. N., Nguyen, A. L., Hoyt, C. R., McIntyre, L., Earl, E. A., & Klein, R. L. (2018). Behavioral interventions for reducing head motion during MRI scans in children. NeuroImage, 171, 234–245.

Greene, D. J., Marek, S., Gordon, E. M., Siegel, J. S., Gratton, C., Laumann, T. O., Gilmore, A. W., Berg, J. J., Nguyen, A. L., & Dierker, D. (2019). Integrative and network-specific connectivity of the basal ganglia and thalamus defined in individuals. Neuron.

Hodgson, K., Poldrack, R. A., Curran, J. E., Knowles, E. E., Mathias, S., Göring, H. H., Yao, N., Olvera, R. L., Fox, P. T., & Almasy, L. (2017). Shared genetic factors influence head motion during MRI and body mass index. Cerebral Cortex, 27(12), 5539–5546.

Kong, R., Li, J., Orban, C., Sabuncu, M. R., Liu, H., Schaefer, A., Sun, N., Zuo, X.-N., Holmes, A. J., & Eickhoff, S. B. (2019). Spatial topography of individual-specific cortical networks predicts human cognition, personality, and emotion. Cerebral Cortex, 29(6), 2533–2551.

Krienen, F. M., Yeo, B. T., & Buckner, R. L. (2014). Reconfigurable task-dependent functional coupling modes cluster around a core functional architecture. Philosophical Transactions of the Royal Society B: Biological Sciences, 369(1653), 20130526.

Laumann, T. O., Gordon, E. M., Adeyemo, B., Snyder, A. Z., Joo, S. J., Chen, M.-Y., Gilmore, A. W., McDermott, K. B., Nelson, S. M., & Dosenbach, N. U. (2015). Functional system and areal organization of a highly sampled individual human brain. Neuron, 87(3), 657–670.

Laumann, T. O., Snyder, A. Z., Mitra, A., Gordon, E. M., Gratton, C., Adeyemo, B., Gilmore, A. W., Nelson, S. M., Berg, J. J., & Greene, D. J. (2016). On the stability of BOLD fMRI correlations. Cerebral Cortex, 27(10), 4719–4732.

Marcus, D., Harwell, J., Olsen, T., Hodge, M., Glasser, M., Prior, F., Jenkinson, M., Laumann, T., Curtiss, S., & Van Essen, D. (2011). Informatics and data mining tools and strategies for the human connectome project. Frontiers in Neuroinformatics, 5, 4.

Marek, S., Siegel, J. S., Gordon, E. M., Raut, R. V., Gratton, C., Newbold, D. J., Ortega, M., Laumann, T. O., Adeyemo, B., & Miller, D. B. (2018). Spatial and temporal organization of the individual human cerebellum. Neuron, 100(4), 977–993.

Miezin, F. M., Maccotta, L., Ollinger, J. M., Petersen, S. E., & Buckner, R. L. (2000). Characterizing the hemodynamic response: Effects of presentation rate, sampling procedure, and the possibility of ordering brain activity based on relative timing. Neuroimage, 11(6), 735–759.

Miranda-Dominguez, O., Mills, B. D., Carpenter, S. D., Grant, K. A., Kroenke, C. D., Nigg, J. T., & Fair, D. A. (2014). Connectotyping: Model based fingerprinting of the functional connectome. PloS One, 9(11), e111048.

Mueller, S., Wang, D., Fox, M. D., Yeo, B. T., Sepulcre, J., Sabuncu, M. R., Shafee, R., Lu, J., & Liu, H. (2013). Individual variability in functional connectivity architecture of the human brain. Neuron, 77(3), 586–595.

Nikolaidis, A., Heinsfeld, A. S., Xu, T., Bellec, P., Vogelstein, J., & Milham, M. (2020). Bagging improves reproducibility of functional parcellation of the human brain. NeuroImage, 116678.

Noble, S., Spann, M. N., Tokoglu, F., Shen, X., Constable, R. T., & Scheinost, D. (2017). Influences on the test–retest reliability of functional connectivity MRI and its relationship with behavioral utility. Cerebral Cortex, 27(11), 5415–5429.

Ojemann, J. G., Akbudak, E., Snyder, A. Z., McKinstry, R. C., Raichle, M. E., & Conturo, T. E. (1997). Anatomic localization and quantitative analysis of gradient refocused echo-planar fMRI susceptibility artifacts. Neuroimage, 6(3), 156–167.

Poldrack, R. A., Laumann, T. O., Koyejo, O., Gregory, B., Hover, A., Chen, M.-Y., Gorgolewski, K. J., Luci, J., Joo, S. J., & Boyd, R. L. (2015). Long-term neural and physiological phenotyping of a single human. Nature Communications, 6(1), 1–15.

Power, J. D., Cohen, A. L., Nelson, S. M., Wig, G. S., Barnes, K. A., Church, J. A., Vogel, A. C., Laumann, T. O., Miezin, F. M., & Schlaggar, B. L. (2011). Functional network organization of the human brain. Neuron, 72(4), 665–678.

Power, J. D., Mitra, A., Laumann, T. O., Snyder, A. Z., Schlaggar, B. L., & Petersen, S. E. (2014). Methods to detect, characterize, and remove motion artifact in resting state fMRI. Neuroimage, 84, 320–341.

Power, J. D., Schlaggar, B. L., Lessov-Schlaggar, C. N., & Petersen, S. E. (2013). Evidence for hubs in human functional brain networks. Neuron, 79(4), 798–813.

Seitzman, B. A., Gratton, C., Laumann, T. O., Gordon, E. M., Adeyemo, B., Dworetsky, A., Kraus, B. T., Gilmore, A. W., Berg, J. J., & Ortega, M. (2019). Trait-like variants in human functional brain networks. Proceedings of the National Academy of Sciences, 116(45), 22851–22861.

Smith, S. M., Jenkinson, M., Woolrich, M. W., Beckmann, C. F., Behrens, T. E., Johansen-Berg, H., Bannister, P. R., De Luca, M., Drobnjak, I., & Flitney, D. E. (2004). Advances in functional and structural MR image analysis and implementation as FSL. Neuroimage, 23, S208–S219.

Sorensen, T. A. (1948). A method of establishing groups of equal amplitude in plant sociology based on similarity of species content and its application to analyses of the vegetation on Danish commons. Biol. Skar., 5, 1–34.

Sylvester, C. M., Yu, Q., Srivastava, A. B., Marek, S., Zheng, A., Alexopoulos, D., Smyser, C. D., Shimony, J. S., Ortega, M., & Dierker, D. L. (2020). Individual-specific functional connectivity of the amygdala: A substrate for precision psychiatry. Proceedings of the National Academy of Sciences, 117(7), 3808– 3818.

Talairach, J. (1988). 3-dimensional proportional system; an approach to cerebral imaging. Co-planar stereotaxic atlas of the human brain. Thieme, 1–122.

Vanderwal, T., Eilbott, J., & Castellanos, F. X. (2019). Movies in the magnet: Naturalistic paradigms in developmental functional neuroimaging. Developmental Cognitive Neuroscience, 36, 100600.

Yeo, B. T., Krienen, F. M., Sepulcre, J., Sabuncu, M. R., Lashkari, D., Hollinshead, M., Roffman, J. L., Smoller, J. W., Zöllei, L., & Polimeni, J. R. (2011). The organization of the human cerebral cortex estimated by intrinsic functional connectivity. Journal of Neurophysiology, 106(3), 1125–1165.

